# Structural characterization of a minimal KLC2/Nup358/BicD2 complex

**DOI:** 10.64898/2026.01.17.700114

**Authors:** Crystal R Noell, Sozanne R Solmaz

## Abstract

Cellular transport processes along microtubules are often facilitated by multi-motor complexes, which are connected by adapter proteins and cargoes. The nuclear pore protein Nup358, for example, interacts with the dynein adapter Bicaudal D2 (BicD2), which in turn recruits minus-end directed dynein motors and plus-end directed kinesin-1 motors for a nuclear positioning pathway that is essential for brain development. How motor recruitment is regulated by interactions of BicD2 with Nup358 is not well understood. Here, we characterize the structure of a minimal complex of kinesin-1 light chain 2 (KLC2), Nup358 and BicD2 by cryo-electron microscopy and small angle X-ray scattering. KLC2/Nup358 assumes a rod-like structure that increases in thickness, when BicD2 is bound. The addition of BicD2 also shifts the KLC2/Nup358/BicD2 complex towards a 2:2:2 stoichiometry, promoting dimerization at lower protein concentrations than without BicD2. Similarly, the presence of the Nup358/KLC2 interaction results in a shift towards a 2:2:2 stoichiometry. Based on these results, we hypothesize that KLC2 and BicD2 are recruited to Nup358 in a cooperative manner, and cooperativity may be promoted by modulation of the oligomeric state.

## INTRODUCTION

In cells, the transport of cargoes along microtubules (MTs) is often facilitated by multi-motor complexes, which are connected by adapter proteins and cargoes (Cason & Holzbaur, 2022; Dharan et al., 2017; Fu & Holzbaur, 2014; Grigoriev et al., 2007; Hendricks et al., 2010; Morris & Hollenbeck, 1993; Wilson & Holzbaur, 2012, 2015). For example, the dynein adapter Bicaudal D2 (BicD2) forms interactions with cargoes and recruits both minus-end directed dynein motors, as well as plus-end directed kinesin-1 motors to the complex. In this arrangement, dynein binds to coiled coil domain 1 of BicD2, kinesin-1 binds to coiled coil domain 2, and cargo binds to coiled coil domain 3 of BicD2 (Grigoriev et al., 2007; Hoogenraad et al., 2001, 2003; Matanis et al., 2002; Schlager, Serra-Marques, et al., 2014; Splinter et al., 2010, 2012). However, the mechanism for motor recruitment to multi-motor complexes is complicated, and the details remain elusive. Dynein adapters such as BicD2 have key roles in cellular transport pathways, as they link the motors to the cargoes and are also required to activate dynein for processive motility (McKenney et al., 2014; Schlager, Hoang, et al., 2014; Splinter et al., 2012; Urnavicius et al., 2015). A cargo for BicD2 is Nup358 (also known as RanBP2), a subunit of the nuclear pore complex. Nup358 interacts with BicD2, which in turn recruits dynein and kinesin-1 motors to facilitate a nuclear positioning pathway that is required for the differentiation of brain progenitor cells, which give rise to most neurons and glia cells in the brain (Hu et al., 2013). The physiological importance of BicD2 in brain development is underscored by the fact that human disease mutations cause devastating neuromuscular and brain developmental diseases, including spinal muscular atrophy, the most common genetic cause of infant death (Huang & Fan, 2017; Neveling et al., 2013; Oates et al., 2013; Peeters et al., 2014; Ravenscroft et al., 2016; Synofzik et al., 2014; Tsai et al., 2020; Yi et al., 2023).

Kinesin-1 is a tetramer that consists of two light chain (KLC) and two heavy chain (KHC) subunits (Vale et al., 1985). KLCs have four isoforms (KLC1–4) in mammals. The KLCs are important for cargo recognition and cargo-driven activation, whereas the KHCs contain the motor domain. Short sequence motifs such as LEWD are sufficient for a cargo to be recognized by kinesin-1 light chains via the TPR domain (tetratricopeptide repeat-containing domain) and for activation of processive motility; the mutation of the LEWD motif to LEAA disrupts the interaction with KLCs (Dodding et al., 2011; Pernigo et al., 2013; Sanger et al., 2017). However, additional interactions between cargo and kinesin-1 are needed to achieve full motor activity, and the mechanistic details of kinesin-1 activation are still debated (Ali et al., 2025; Blasius et al., 2007; Chiba et al., 2022; Cockburn et al., 2018; Cross et al., 2024; Dimitrova-Paternoga et al., 2021; Heber et al., 2024; Sanger et al., 2017; Shukla et al., 2025; Tan et al., 2023; Weijman et al., 2026; Yip et al., 2016a).

Previously, we established a structural model for a minimal complex of Nup358 and BicD2 by structure prediction with AlphaFold, followed by experimental validation of the model by NMR, mutagenesis, binding assays, and functional assays (Fig. 1A) (Gibson et al., 2022, 2023). In the complex, the BicD2 binding site is separated by a ∼30 residue flexible linker from the LEWD sequence motif of Nup358 that acts as kinesin-1 binding site (Gibson et al., 2022, 2023). To confirm that BicD2 and KLC2 interact simultaneously with Nup358, we assembled a minimal KLC2/Nup358/BicD2 complex and established that it has a 2:2:2 stoichiometry (Cui et al., 2019). To further confirm these results, we created a fusion protein of Nup358 (human Nup358-min, aa 2148-2240) and fused it to the N-terminus of KLC2 (Fig. 1C, mouse, aa 196-480) and then investigated the interaction of this Nup358-KLC2 fusion protein with the human BicD2 cargo-binding domain (BicD2-CTD, aa 715-804) (Cui et al., 2019). The design of this fusion protein was based on a previous study, where the LEWD motif of the protein SKIP (SifA-kinesin interacting protein) was fused to KLC2 with a short ∼16 aa flexible linker. A crystal structure confirmed that the LEWD motif interacts with KLC2 within the fusion protein (Fig. 1B) (Pernigo et al., 2013).

**Figure 1.**
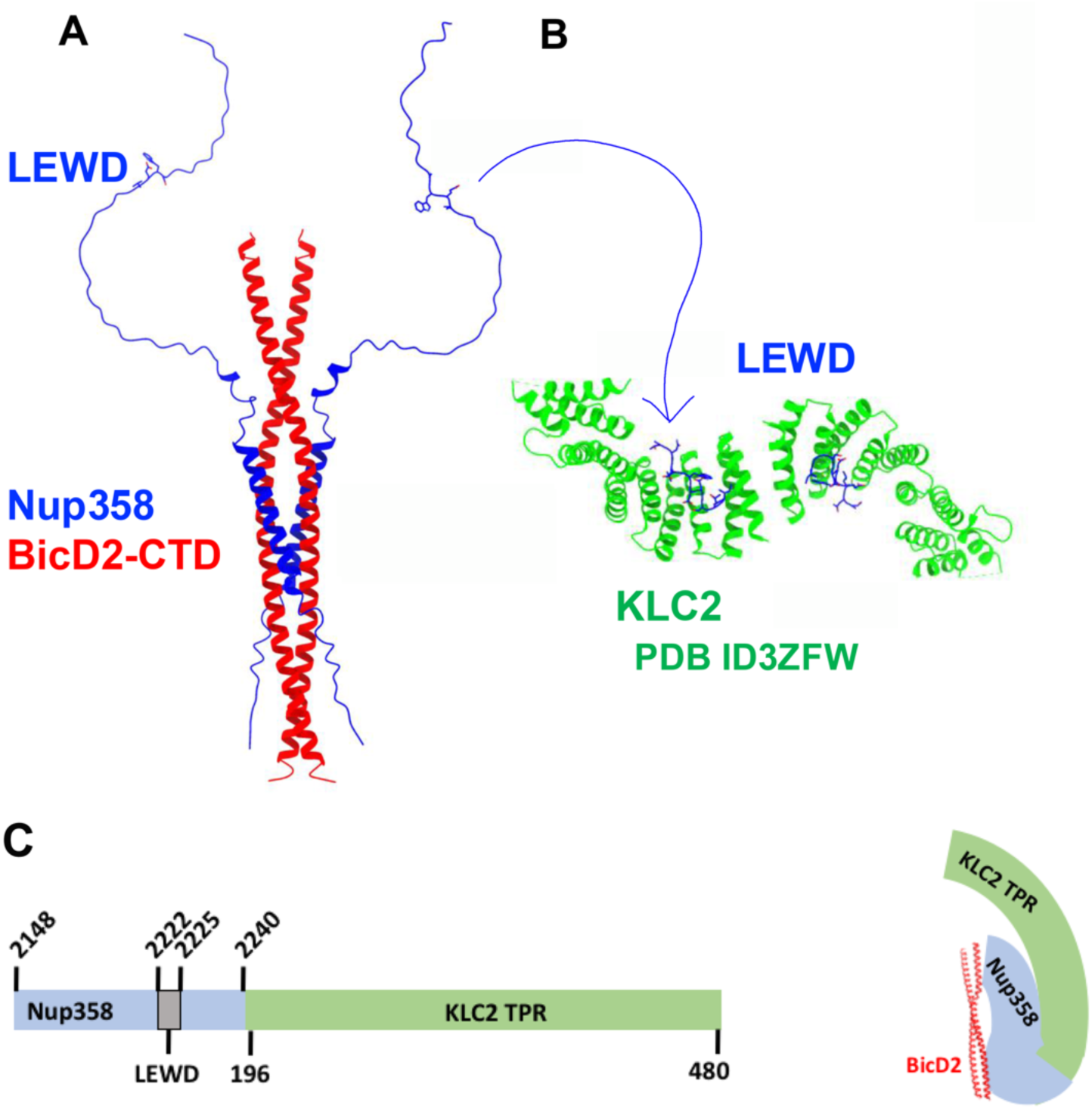
A minimal complex from KLC2, Nup358 and BicD2 was previously assembled which has a 2:2:2 stoichiometry. (A) The experimentally validated structural model of the minimal Nup358/BicD2 complex is shown in cartoon representation; the KLC2 recruiting LEWD motif is indicated and shown in bonds representation (Gibson et al., 2022, 2023). (B) Cartoon representation of the X-ray structure of a fusion protein of the TPR domain of KLC2 (green) and a peptide with a LEWD motif (blue) (Pernigo et al., 2013). (C) A schematic representation of the Nup358-KLC2 fusion protein and its complex with BicD2-CTD (Cui et al., 2019).

Similar to a complex assembled from the individual proteins, the Nup358-KLC2 fusion protein dimerizes and also interacts with the BicD2-CTD to form a 2:2 complex, suggesting that the LEWD motif of Nup358 forms an intramolecular interaction with KLC2 within the fusion protein (Cui et al., 2019). The fusion protein offers the advantage of being a single peptide, eliminating the formation of a variety of oligomeric states. The fusion also locally increases the concentration of the interacting partners, thereby strengthening the interaction (Cui et al., 2019). We have also previously shown that mutation of the LEWD motif of Nup358 to the sequence LEAA virtually abolished the interaction with the KLC2 (Cui et al., 2019). A W2224A/D2225A variant of the Nup358-KLC2 fusion protein forms predominantly monomers, whereas the WT complex dimerizes, suggesting that dimerization is promoted by the intramolecular interaction of the LEWD motif of Nup358 with KLC2 (Cui et al., 2019). A general hypothesis for kinesin-1 activation suggests that interaction of the kinesin-1 light chains with cargoes promotes dimerization of the light chains, which is one of the requirements for cargo-driven activation of kinesin-1 motors (Cockburn et al., 2018). In line with this hypothesis, our previous results suggest that the interaction of Nup358 and KLC2 results in 2:2 complex formation, and the W-acidic LEWD motif is required for the oligomerization, while KLC2 and Nup358 form predominantly monomers (Cui et al., 2019).

Here we use single-particle cryo-electron microscopy and small-angle X-ray scattering to structurally characterize a minimal complex of KLC2, Nup358 and BicD2. Both the KLC2/Nup358 complex and the ternary complex with BicD2 form rod-like structures, and the thickness of the rod increases when BicD2 is bound. Furthermore, addition of BicD2 to Nup358/KLC2 promotes oligomerization, which may facilitate cooperative recruitment of KLC2 and BicD2 to Nup358.

## RESULTS

### Structural characterization of a minimal KLC2/Nup358/BicD2 complex by single-particle cryo-EM

In order to perform a structural characterization of the interactions between KLC2, BicD2 and Nup358, we assembled a complex of the Nup358-KLC2 fusion protein (Nup358, aa 2148-2240 is fused to the N-terminus of KLC2, aa 196-480) and the human BicD2-CTD (cargo-binding domain, aa 715-804) (Cui et al., 2019) and analyzed it by single-particle cryo-electron microscopy (cryo-EM). This complex is further described in the introduction and in Fig. 1. Fig. 2 shows a representative cryo-electron micrograph and representative 2D classes of 1,355,298 particles from 4188 micrographs. The 3D reconstruction is shown in Fig. 3, which suggests that the complex has an S-like shape. Structural details like the locations of the alpha-helices of the TPR domain of KLC2 are resolved in the complex. The KLC2-TPR domain is well-defined (Fig. 1) but does not account for all of the observed electron density in the map. Further improvement of the structure is in progress in order to obtain more structural information for bound Nup358 and BicD2. We conclude that the S-shaped complex includes dimers of KLC2 and in addition some domains of Nup358 and BicD2.

**Fig. 2.**
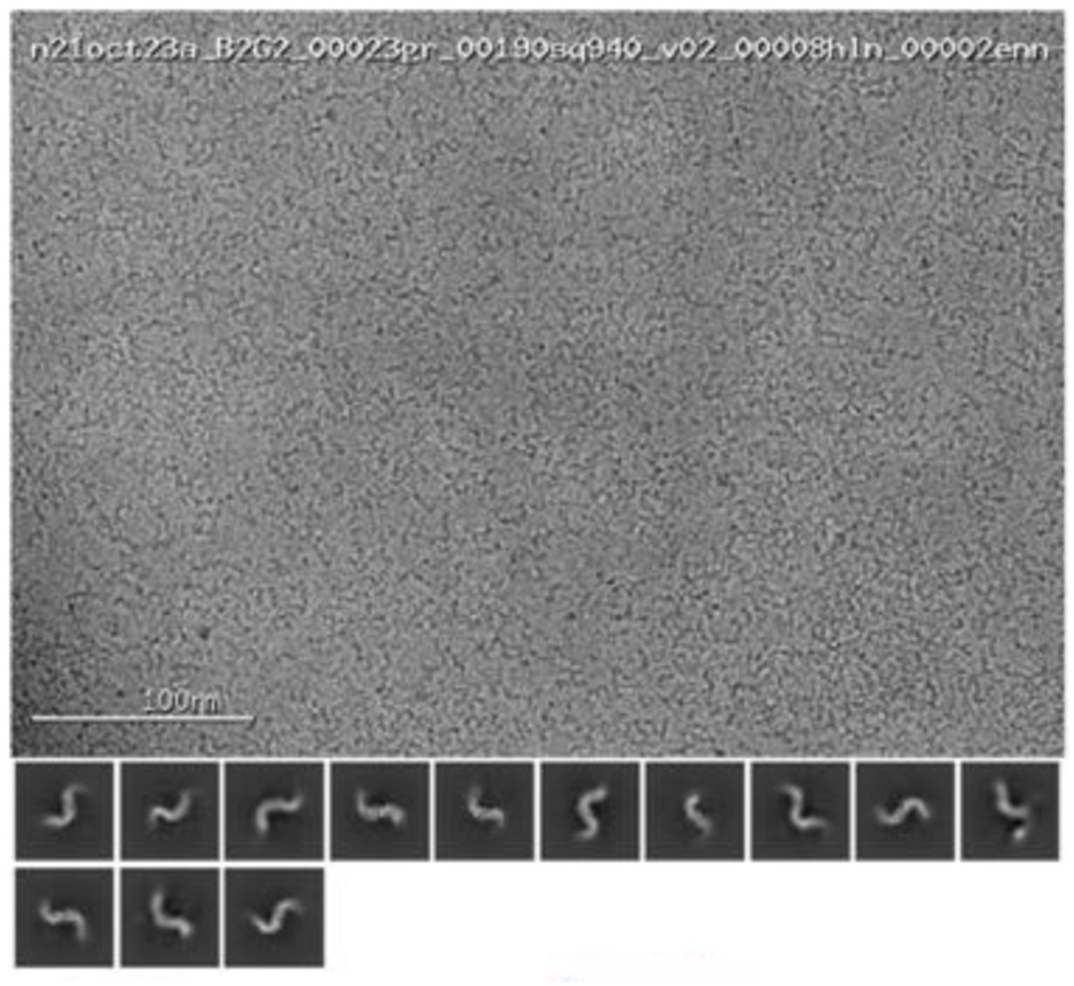
Structural analysis of the minimal Nup358-KLC2/BicD2-CTD complex by single-particle cryo-EM. Representative cryo-electron micrograph and 2D classes of 1,355,298 particles from 4188 micrographs.

**Fig. 3.**
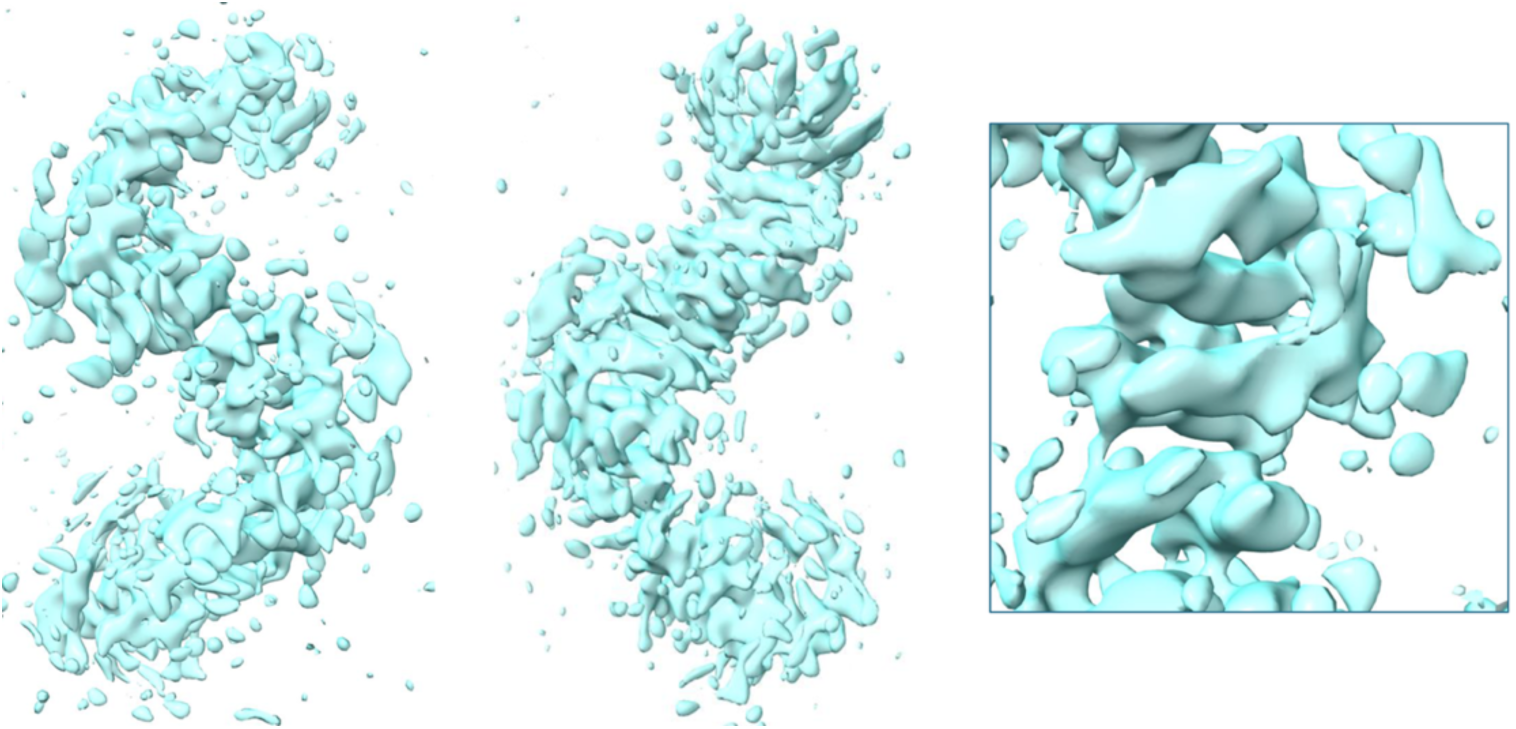
Structural analysis of the minimal Nup358-KLC2/BicD2 complex by single-particle cryo-EM. 3D reconstruction is shown. GSFC resolution= 5.2 Å.

### Both BicD2 and KLC2 promote oligomerization with Nup358

To investigate the structural relationship between KLC2, Nup358 and BicD2 within the triple complex, we analyzed the Nup358-KLC2 fusion protein and the complex with BicD2-CTD added using solution Small-angle X-ray scattering (SAXS). One difference of SAXS compared to cryo-EM is the fact that intrinsically disordered regions contribute to the overall molecular envelope established by SAXS, whereas they are typically averaged out in cryo-EM due to conformational flexibility. SAXS scattering profiles were collected from a range of protein concentrations (Table 1, Fig. 2, Figs. S1 - S30), in order to determine the concentration dependence of the molar mass and of the radius of gyration (R_G_) in solution. R_G_ is the radius of a hollow sphere which would have the same moment of inertia as the actual body, if the entire mass of the body was located there (Alsaker et al., 2019). It gives an idea of the overall shape of a molecule (Hopkins et al., 2017).

**Table 1.** Guinier and P(r) analysis by SAXS.

| Concentration<br>(mg/mL) | R<sub>G</sub> (Å)<br>Guinier<br>plot | R<sub>G</sub> (Å)<br>P(r)<br>function | I(0)<br>P(r) function | TE / $\chi^2$<br>P(r)<br>function | D<sub>Max</sub><br>P(r)<br>function | MW(kDa)<br>I(0)<sub>ref</sub> /<br>I(0)<sub>abs</sub><br>(± 10%<br>error) | OS |
| --- | --- | --- | --- | --- | --- | --- | --- |
| **Nup358-KLC2 Fusion protein** |  |  |  |  |  |  |  |
| 3.0 | 41.0 ±0.1 | 47.6 ±0.6 | 0.164 ±0.002 | 0.73/ 1.05 | 180 | 76.4/ 67.9 | 1.8 |
| 2.0 | 39.4 ±0.2 | 45.4 ±0.6 | 0.100 ±0.001 | 0.72/ 1.09 | 180 | 70.0/ 62.1 | 1.6 |
| 1.6 | 40.2 ±0.2 | 44.8 ±0.7 | 0.0802 ±0.001 | 0.62/ 1.04 | 170 | 73.0/ 64.8 | 1.7 |
| 1.2 | 38.9 ±0.3 | 45.5 ±0.5 | 0.0589 ±0.0006 | 0.73/ 1.23 | 175 | 68.3/ 60.6 | 1.6 |
| 1.0 | 40.5 ±0.2 | 44.6 ±0.4 | 0.0409 ±0.0003 | 0.69/ 1.22 | 175 | 61.0/ 54.2 | 1.4 |
| 0.8 | 39.7 ±0.2 | 41.4 ±0.2 | 0.0366 ±0.0007 | 0.70/ 1.05 | 175 | 70.1/ 62.3 | 1.6 |
| 0.6 | 39.9 ±0.3 | 44.8 ±0.8 | 0.0263 ±0.0004 | 0.70/ 1.13 | 175 | 64.7/ 57.5 | 1.5 |
| 0.4 | 39.8 ±0.5 | 45.8 ±1.4 | 0.0168 ±0.0004 | 0.57/ 1.08 | 175 | 60.5/ 53.8 | 1.4 |
| 0.2 | 37.4 ±0.8 | 41.9 ±1.6 | 0.0061 ±0.0002 | 0.61/ 1.09 | 150 | 44.5/ 39.5 | 1.0 |
| **Nup358-KLC2 fusion protein + BicD2-CTD 1:1 molar ratio** |  |  |  |  |  |  |  |
| 3.75 | 44.4 ±0.2 | 50.6 ±0.4 | 0.2791 ±0.002 | 0.80/ 1.09 | 190 | 105.1/ 93.3 | 2.0 |
| 2.5 | 43.8 ±0.2 | 50.0 ±0.4 | 0.2016 ±0.002 | 0.79/ 1.14 | 190 | 113.8/ 101.1 | 2.1 |
| 2.0 | 44.8 ±0.2 | 52.1 ±0.3 | 0.1635 ±0.001 | 0.75/ 1.38 | 200 | 115.2/ 102.3 | 2.1 |
| 1.5 | 45.5 ±0.2 | 49.9 ±0.4 | 0.1116 ±0.0006 | 0.70/ 1.13 | 191 | 110.9/ 98.5 | 2.1 |
| 1.2 | 43.6 ±0.2 | 48.9 ±0.6 | 0.0899 ±0.0009 | 0.69/ 1.04 | 190 | 104.9/ 93.1 | 2.0 |
| 1.0 | 44.9 ±0.4 | 49.1 ±0.5 | 0.0720 ±0.0005 | 0.73/ 1.09 | 190 | 107.8/ 95.7 | 2.0 |
| 0.7 | 40.4 ±0.2 | 49.5 ±1.4 | 0.0529 ±0.0002 | 0.69/ 1.27 | 190 | 93.0/ 82.6 | 1.7 |
| 0.5 | 40.9 ±0.2 | 48.6 ±1.0 | 0.0319 ±0.0005 | 0.72/ 1.45 | 190 | 88.4/ 78.5 | 1.6 |
| 0.2 | 40.3 ±0.3 | 48.2 ±1.4 | 0.0140 ±0.0003 | 0.60/ 1.39 | 185 | 79.2/ 70.4 | 1.5 |
| **Nup358/W2224A/D2225A-KLC2 fusion protein** |  |  |  |  |  |  |  |
| 2.0 | 37.5 ±0.4 | 41.2 ±0.6 | 0.0795 ±0.0006 | 0.65/ 1.01 | 175 | 60.6/ 53.8 | 1.4 |
| 1.6 | 37.9 ±0.4 | 41.6 ±0.7 | 0.0601 ±0.0005 | 0.66/ 1.11 | 175 | 57.1/ 50.8 | 1.3 |
| 1.2 | 39.6 ±0.4 | 43.4 ±0.7 | 0.0539 ±0.0005 | 0.69/ 1.06 | 175 | 68.0/ 60.4 | 1.6 |
| 1.0 | 37.4 ±0.1 | 41.2 ±0.6 | 0.0382 ±0.0003 | 0.65/ 1.03 | 175 | 58.2/ 51.7 | 1.4 |
| 0.8 | 36.3 ±0.3 | 40.3 ±0.5 | 0.0300 ±0.0002 | 0.57/ 1.16 | 160 | 56.9/ 50.5 | 1.3 |
| 0.6 | 36.0 ±0.4 | 38.7 ±0.8 | 0.0222 ±0.0002 | 0.63/ 1.12 | 152 | 56.9/ 50.5 | 1.3 |
| 0.4 | 34.4 ±0.6 | 39.5 ±1.0 | 0.0138 ±0.0002 | 0.64/ 1.09 | 157 | 51.5/ 45.8 | 1.2 |
| **Nup358/W2224A/D2225A-KLC2 fusion protein + BicD2-CTD in 1:1 ratio** |  |  |  |  |  |  |  |
| 1.5 | 44.4 ±0.6 | 46.3 ±0.8 | 0.0962 ±0.001 | 0.65/ 0.99 | 180 | 100.2/ 89.0 | 1.9 |
| 1.25 | 43.6 ±0.5 | 48.5 ±0.7 | 0.0839 ±0.0008 | 0.64/ 1.06 | 190 | 101.8/ 90.4 | 1.9 |
| 1.0 | 41.4 ±0.5 | 46.7 ±0.7 | 0.0594 ±0.0006 | 0.60/ 1.06 | 190 | 89.3/ 79.3 | 1.7 |
| 0.75 | 42.5 ±0.5 | 47.1 ±0.6 | 0.0448 ±0.0004 | 0.54/ 1.00 | 185 | 90.9/ 80.7 | 1.7 |
| 0.5 | 37.4 ±0.4 | 39.2 ±0.7 | 0.0252 ±0.0003 | 0.61/ 1.08 | 150 | 78.4/ 69.6 | 1.5 |
| 0.25 | 32.0 ±0.7 | 37.0 ±2.8 | 0.0115 ±0.0004 | 0.61/ 1.11 | 159 | 68.9/ 61.2 | 1.3 |

R_G_: radius of gyration. I(0): scattering intensity at zero angle. TE: total estimate. D_Max_: maximum particle diameter. MW: molar mass. OS: Calculated oligomeric state, MW/monomer mass. Molar masses were determined with the I(0) ref method (Mylonas 2007) and an additional molecular mass using I(0)_abs_ method is displayed (Jeffries et al., 2016). For additional SAXS analyses, see Fig. S1 - S30 and Table S1.

**Figure 4.**
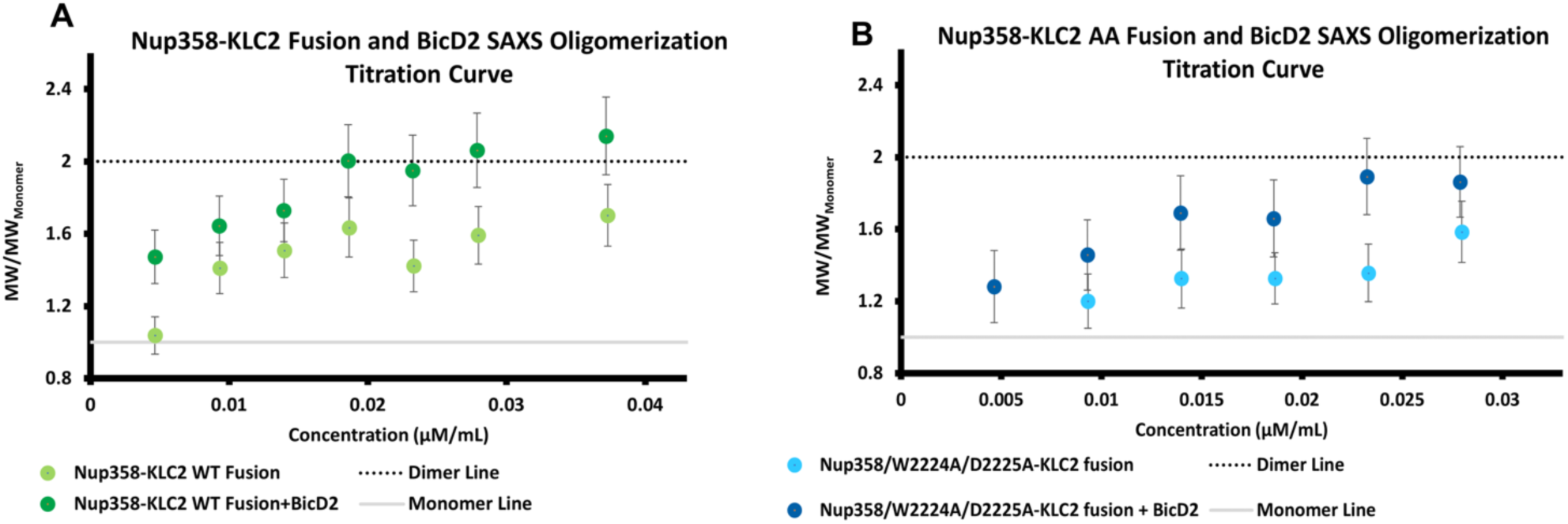
SAXS data indicate that the addition of BicD2 leads to increased oligomerization. (A, B). The oligomeric state is plotted versus the protein concentration. To obtain the oligomeric state, the molar mass from SAXS was divided by the calculated molecular weight of the KLC2-fusion protein (42.9 kDa) or the complex with BicD2-CTD (42.9 kDa + 10.9 kDa). (A) Oligomerization curves of the Nup358-KLC2 WT fusion protein (light green) and the complex with BicD2-CTD (green). (B) Oligomerization curves of the Nup358 W2224A/D2225A-KLC2 fusion protein (light blue) and of the complex with BicD2-CTD (blue). For additional SAXS analyses, see Fig. S1 - S30 and Table S1.

To create a titration curve of the oligomeric state versus the protein concentration, the oligomeric state was obtained by dividing the molar masses from SAXS by the calculated molecular weight of a Nup358-KLC2 fusion protomer (42.9 kDa) or of the complex with BicD2-CTD (53.8 kDa) (Fig. 4). We observed that the addition of BicD2-CTD to the Nup358-KLC2 fusion protein shifted the equilibrium towards 2:2 complex formation compared to the fusion protein alone. The Nup358-KLC2 fusion protein with BicD2-CTD forms a 2:2 complex already at a concentration of ≥ 0.02 µM/mL while the Nup358-KLC2 fusion alone only approaches the dimeric state at concentrations of ≥ 0.04 µM/mL (Fig. 4).

Furthermore, the WT Nup358-KLC2 fusion protein consistently has a lower radius of gyration than the samples with BicD2-CTD added. This size increase is expected with the addition of BicD2-CTD, since it promotes oligomerization.

To assess the impact of the intramolecular interaction between the LEWD motif of Nup358 and KLC2 within the fusion protein, we also performed SAXS experiments with the W2224A/D2225A mutant, in which this interaction is virtually abolished. Molar masses for the Nup358-KLC2 fusion protein determined by SAXS at different protein concentrations range from approximately 60 kDa - 76 kDa (corresponding to an oligomeric state of 1.4 to 1.8). To compare, molar masses of the W2224A/D2225A mutant from SAXS ranged from 51 kDa - 68 kDa, which matches to an oligomeric state of 1.2 to 1.6. For the mixture of the Nup358-KLC2 fusion protein and the BicD2-CTD, molar masses determined by SAXS range from 79 – 115 kDa, which corresponds to an oligomeric state of 1.2 to 2.1. Lastly, molar masses of the fusion W2224A/D2225A mutant protein/BicD2-CTD complex determined by SAXS ranged from 68-101 kDa, which corresponds to an oligomeric state of 1.3 to 1.9.

The Nup358-KLC2/W2224A/D2225A fusion protein/BicD2-CTD complex forms a 2:2 complex at a concentration ≥ 0.025 µM/mL, which is a higher concentration compared to the WT (Fig. 4), suggesting that the Nup358/KLC2 interaction shifts the equilibrium further towards 2:2 complex formation. Furthermore, the Nup358-KLC2/W2224A/D2225A fusion protein without BicD2-CTD only approaches an oligomeric state of 1.6 at the highest concentration that was assessed (0.03 µM/mL) (Fig. 4) indicating that the interaction between KLC2 and Nup358 also shifts the equilibrium to 2:2 complex formation.

To conclude, our oligomerization plots suggest that addition of BicD2-CTD shifts the equilibrium of the Nup358-KLC2 fusion protein towards 2:2 complex formation, and the Nup358/KLC2 interaction also pushes the equilibrium towards 2:2 complex formation.

### The KLC2-Nup358 fusion protein forms a rod-like structure that gets thicker upon addition of BicD2-CTD

To obtain structural information, pair distance distribution (P(r)) functions were determined from the SAXS profiles. The P(r) function is a histogram showing the distribution of the distances between all atom pairs in the structure (Choi & Morais, 2014). Based on the concentration-dependent oligomerization curves determined from SAXS (Fig. 4), we selected protein concentrations for further analysis for which the oligomeric state was as close to a dimer as possible. Comparing the P(r) functions of different samples at similar oligomeric states allows us to identify structural changes that are specific to the addition of BicD2-CTD. The P(r) functions of the Nup358-KLC2 WT fusion protein have a peak that decays with a linear slope, which is typical for a rod-like structure. The thickness of a rod-like structure is similar throughout the rod, therefore the pair distances will have a linear decay, whereas a globular structure would have a gaussian-shaped peak. The main features of the P(r) function are a primary peak at 30 Å and a shoulder peak at 60 Å (labeled in Fig. 5).

**Figure 5.**
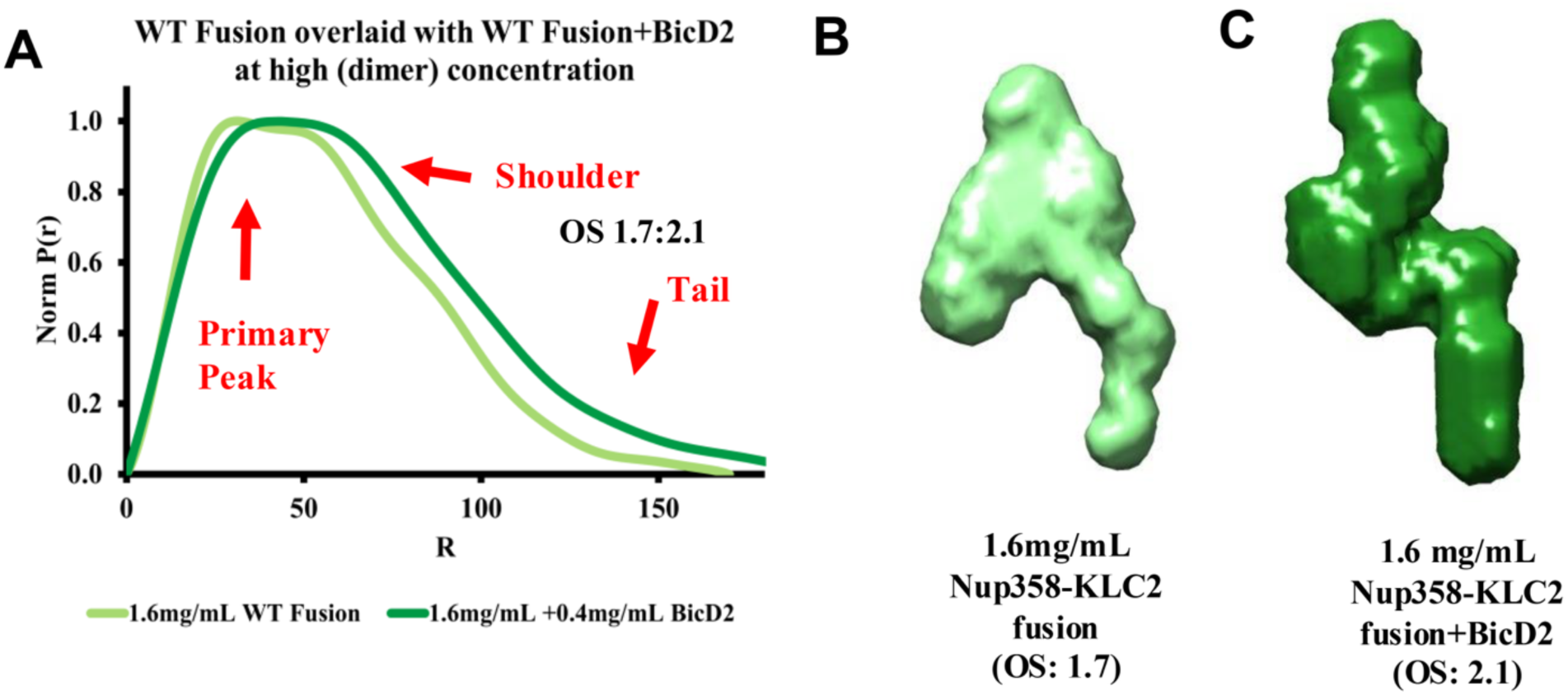
The KLC2-Nup358 fusion protein forms a rod-like structure that thickens upon addition of BicD2-CTD. Solution SAXS data was collected for the Nup358-KLC2 fusion protein as well as the complex with BicD2-CTD. The oligomeric state ratio (OS) is indicated. (A) Normalized P(r) functions of Nup358-KLC2 fusion protein/BicD2-CTD complex (1.6 mg/mL +0.4 mg/mL) (oligomeric state = 2.1; light green) and the Nup358-KLC2 WT fusion protein at 1.6 mg/mL (green; oligomeric state = 1.7). The primary peak at 30 Å, the peak shoulder at 60 Å and the trailing tail from 100 - 200 Å are indicated. (B) Bead model reconstructions of molecular envelopes of the Nup358-KLC2 fusion protein (1.6 mg/mL) and of the Nup358-KLC2 fusion protein/BicD2-CTD complex (1.6 mg/mL). For additional SAXS analyses, see Fig. S1 - S30 and Table S1.

The wild-type Nup358-KLC2 fusion protein /BicD2-CTD complex also has a rod-like shape since the P(r) function has a characteristic linear decay of the main peak. The structure likely also contains a second larger domain to account for the second shoulder peak seen around 60 Å. When the P(r) functions of the Nup358-KLC2 WT fusion proteins are compared with the ones of complexes that contain also BicD2-CTD, it becomes evident that addition of the BicD2-CTD increases the thickness of the rod shape (Fig. 5). In the complex with BicD2-CTD, the linear slope of the decaying peak has shifted towards distances that are approximately 10 Å larger compared to the P(r) functions of the Nup358-KLC2 WT fusion proteins, and the largest diameter D_max_ is slightly larger as well, indicating also an increase of the length of the rod structure.

We also performed bead model reconstructions of the molecular envelopes from our SAXS data (Franke & Svergun, 2009); most of them had NSD (normalized spatial discrepancy) values of around 1 which suggests high structural homogeneity (Table S1). The majority of our samples had a χ^2^ value around one indicating a good fit between the model and the scattering data.

Overall, the molecular envelopes are consistent with the conclusion drawn from the P(r) functions. Both the Nup358/KLC2 fusion protein and the complex with BicD2-CTD added had rod-like structures and addition of BicD2-CTD resulted in a thicker and slightly longer rod structure.

To conclude, our SAXS results indicate that simultaneous interaction of KLC2 and BicD2 with Nup358, shifts the equilibrium towards higher order oligomerization. Furthermore, the KLC2-Nup358 complex forms a rod-like structure which becomes thicker upon addition of BicD2-CTD (Fig. 5).

#### 3.2.7 Absence of the internal interaction between Nup358-min and KLC2 results in structural changes

In the Nup358-KLC2 WT, an internal interaction between the LEWD motif of Nup358 and KLC2 is formed, which is strongly diminished by the W2224A/D2225A mutation (Cui et al., 2019).

Thus, we compared the structure of the WT and W2224A/D2225A mutant of the Nup358-KLC2 fusion protein/BicD2-CTD complex at a concentration of 1.5 mg/mL, where both the WT and mutant form 2:2 complexes. Interestingly, we were able to see large changes in the P(r) function of the W2224A/D2225A mutant (Fig. 6), which has a broad peak at 30 Å with a very small shoulder at 60 Å and a decaying tail at (100-200 Å), strongly characteristic of a rod like shaped structure. To compare, the P(r) function of the WT complex has a much larger shoulder peak at 60 Å, equal in height to the 30 Å peak which could indicate the presence of an additional domain within the elongated structure of the WT (Fig. 6). Bead model reconstructions of the molecular envelope of the wild-type and W2224A/D2225A mutant Nup358-KLC2 fusion protein/BicD2-CTD complexes show structural differences that may be attributed to the strong diminishment of the intramolecular Nup358/KLC2 interaction in the mutant. The wild-type Nup358-KLC2 fusion protein/BicD2-CTD complex forms a rod-like structure with a thicker middle domain (Fig. 6). While the W2224A/D2225A mutant complex has an elongated shape, similar to the wild type, its middle domain appears on average smaller (Fig. 6). Furthermore, upon comparison of the two P(r) functions of the WT and mutant complexes (Fig. 6), the linear decay is shifted in the WT towards larger distances (∼10 Å) compared to the mutant, indicating that the rod shape of the wild-type Nup358-KLC2 fusion protein/BicD2-CTD complex is thicker compared to the W2224A/D2225A mutant (Fig. 6).

**Figure 6.**
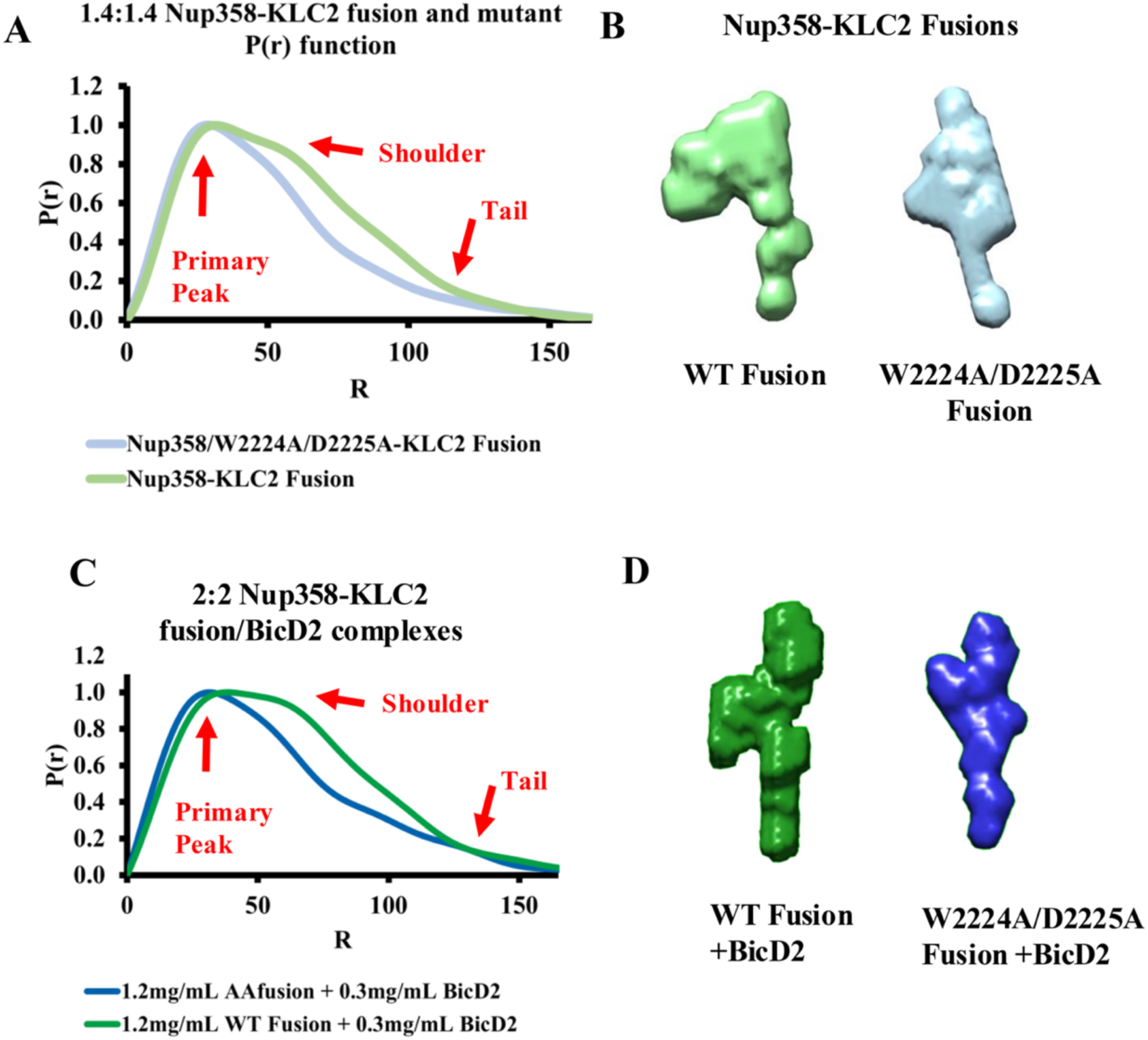
SAXS results provide structural information for the Nup358/KLC2-fusion protein and the W2224A/D2225A mutant, as well as for the ternary complex with BicD2-CTD bound. (A) The P(r) function for the Nup358-KLC2 WT fusion protein (light green) at a concentration of 1 mg/mL and an oligomeric state of 1.4 is shown, together with the P(r) function of the W2224A/D2225A mutant (light blue), at a concentration of 2 mg/mL and an oligomeric state of 1.4. The primary peak at 30 Å, the shoulder at 60 Å and the tail at 100 - 200 Å are labelled in red. (B) Bead model reconstruction of molecular envelopes from the P(r) functions shown in (A). (C) P(r) functions for the Nup358-KLC2-fusion protein/BicD2-CTD complex (green) and the W2224A/D2225A mutant (blue), at a concentration (1.5 mg/ml) where both form 2:2 complexes. (D) Bead model reconstruction of molecular envelopes from the P(r) functions shown in (C). Note that the WT complex (green) shows a mostly symmetrical structure with a larger domain in the middle when compared to the W2224A/D2225A-mutant (blue). For additional SAXS analyses, see Fig. S1 - S30 and Table S1.

Similarly, when comparing P(r) functions for Nup358-KLC2 fusion (1 mg/mL) and the mutant Nup358/W2224A/D2225A-KLC2 fusion (2 mg/mL) proteins in the absence of BicD2-CTD at the same oligomeric state (OS: 1.4) we see structural differences (Fig. 6). The Nup358/W2224A/D2225A-KLC2 fusion protein forms a rod-like structure, but the WT also contains a second domain (at 60 Å) of similar size to the primary rod peak (30 Å). A comparison of the P(r) functions (Fig. 5 and 6) suggests that this second domain peak at 60 Å is due to the Nup358-KLC2 interaction and not due to the addition of BicD2. Instead, the addition of BicD2-CTD appears to show up in the P(r) function as a thickening of the overall peak noticeable between 75 - 125 Å (Fig. 5), indicating it leads to a thickening of the structure rather than specifically increasing the size of the secondary domain with the peak at 60 Å. Furthermore, the D_Max_ of the structures (Fig. 5,6 Table 1) are very similar and the P(r) functions indicate a thicker structure in the presence of BicD2. These SAXS results provide structural insights into the interactions of BicD2-CTD, KLC2 and Nup358-min in the triple complex, and suggest that disruption of the interaction between Nup358 and KLC2 results in structural changes of the triple complex.

## DISCUSSION

Here, we structurally characterize a minimal KLC2/Nup358/BicD2 complex. The 3D reconstruction of the complex from cryo-EM reveals an S-like shape, while complementary SAXS analysis was also performed to visualize additional flexible regions that may be averaged out in cryo-EM. The complex assumes an extended rod-like structure both with and without BicD2 bound, however, the addition of BicD2 increases the thickness of the rod. We also show that addition of BicD2-CTD to the Nup358-KLC2 fusion protein shifts the oligomeric state towards a 2:2 complex, whereas a higher protein concentration is needed to dimerize the Nup358-KLC2 fusion protein in absence of BicD2. Similarly, in the presence of the KLC2/Nup358 interaction, formation of a 2:2 complex is also increased. If the interaction is diminished by a mutation, a much higher protein concentration is required to reach a 2:2 stoichiometry, highlighting the importance of the LEWD motif for cargo-mediated dimerization.

Based on these results, we hypothesize that the recruitment of KLC2 and BicD2 to Nup358 is cooperative, favoring formation of a ternary complex with both proteins bound. This cooperativity may be facilitated by modulation of the oligomeric state. Several recent studies suggest that the oligomeric state is a key regulatory mechanism in many cellular transport pathways (Chaaban S & Carter AP, 2022; McClintock et al., 2018; Sladewski et al., 2018; Urnavicius et al., 2018). Although KLC2 is monomeric in isolation (Cui et al., 2019; Nguyen et al., 2017), it forms 2:2 complexes in the presence of cargo, and the transition may be essential for activation (Cockburn et al., 2018; Nguyen et al., 2017; Yip et al., 2016b; Zhu et al., 2012), as KLC2 also forms dimers within active kinesin-1 motors. Future studies are needed to confirm cooperative recruitment of BicD2 and KLC2 to Nup358 and to establish the specific role of the oligomeric state in promoting such cooperativity.

## METHODS

### Single particle cryo-electron microscopy

The Nup358-KLC2 fusion protein and the BicD2-CTD were expressed and purified as described (Cui et al., 2019). The ternary complex was assembled by mixing these two proteins in 1:1 molar ratio. Frozen sample grids were prepared with the TFS Vitrobot Mark IV plunge-freezer system, by applying 3 µl of the ternary complex with a blot time of 2 s to plasma cleaned holey carbon grids. Images were collected with the Leginon software (Carragher et al., 2000) at NCCAT, using a TFS Krios TEM operating at 300 kV and a Gatan K3 direct electron detector for data collection.

The pixel size was 0.825 Å and the total exposure dose was 60 e^-^/Å^2^. CTF estimation was performed by CTFFind4 (Rohou & Grigorieff, 2015). Particles were selected using the program Topaz (Bepler et al., 2019) and particle extraction and 2D classification was performed in CryoSPARC (Punjani et al., 2017). Ab-initio reconstruction and 3D refinement was also performed in CryoSPARC (Punjani et al., 2017).

### Solution Small angle X-ray scattering (SAXS)

Proteins were expressed and purified as described (Cui et al., 2019). The homogeneity and monodispersity of our samples was confirmed by the SEC-MALS experiments (Cui et al., 2019). The samples were filtered (0.2 µM pore size) and centrifuged for 30 minutes at 4°C at 21,700 g within one to three hours before data collection. For each sample, the SAXS profile of the corresponding buffer match was collected, which was the flow through from the last protein concentration step. SAXS data was collected at Cornell University High Energy Synchrotron Source at the BioSAXS beamline 1D7A1 with a MacCHESS EIGER 4M detector (Dectris) at a single position. A quartz capillary with a 1.48 mm path length was used as the sample cell with OD of 1.5 mm and wall thickness of 10 µm. For each SAXS diffraction profile, 20 diffraction images were collected with an exposure time of 0.5 s and 10-fold attenuation of the beam. The beam current was 100 mA, and data was collected at a temperature of 4°C. The wavelength was 10.02538 keV, the beam dimensions were 0.25 mm x 0.25 mm and the beam flux was 2×10^12^. Twenty SAXS diffraction frames were collected for each sample, and the sample cell was cleaned after each run. SAXS profiles of glucose-isomerase were collected as standards for MW determination. Upon averaging the 20 frames, our samples showed no signs of radiation damage. SAXS diffraction profiles were analyzed using the BioXTAS RAW software (Hopkins et al., 2017; Skou et al., 2014) as well as ATSAS extensions (Manalastas-Cantos et al., 2021).

Twenty SAXS diffraction images were averaged for each sample and for each buffer match. However scattering profiles that were not similar enough to be averaged were excluded. The averaged buffer match SAXS profile was subtracted from the sample SAXS profile. This resulting subtracted file was used for further data analysis.

A Guinier fit (Guinier & Fournet, 1947) was then performed on the subtracted scattering profiles. For molar mass determination, the RAW and ATSAS packages were used (Hopkins et al., 2017; Konarev et al., 2003). The molar mass was calculated with the I(0)ref method (Mylonas & Svergun, 2007) and the I(0)_abs_ method (Orthaber et al., 2000). GNOM (Svergun, 1992), which is implemented in the ATSAS package (Manalastas-Cantos et al., 2021) was used to generate P(r) plots (Svergun, 1992). 15 DAMMIF bead model reconstructions of the molecular envelope were carried out for each sample in ATSAS implemented in RAW (Hopkins et al., 2017; Konarev et al., 2003). Shape reconstructions were generated by DAMMIF/N, an average consensus shape is made using program DAMAVER. Finally DAMCLUST alignment was used to cluster the models into groups of similar models (Franke & Svergun, 2009; Kelley et al., 1996; Svergun, 1999). The models were averaged and refined. Figures of the bead models were created in Chimera (Pettersen et al., 2021).

## ACKNOWLEDGEMENTS

SR Solmaz was supported by NIH grants R01 GM144578 and R15 GM128119. Cryo-EM data was collected at the National Center for CryoEM Access and Training (NCCAT) and the Simons Electron Microscopy Center located at the New York Structural Biology Center, supported by the NIH Common Fund Transformative High Resolution Cryo-Electron Microscopy program (U24 GM129539, and NIGMS R24 GM154192) and by grants from the Simons Foundation (SF349247) and NY State Assembly. We thank NCCAT staff scientists Dr. Eugene Chua and Dr. Huihui Kuang for valuable discussions. SAXS data was collected at beamline 7A1, Cornell High Energy Synchrotron Source, supported by NSF award DMR-1829070, and by NIH/NIGMS award GM-124166. We thank Qingqiu Huang and Richard Gillilan for user support at the synchrotron source. The authors declare that they have no conflict of interest.

## SUPPORTING INFORMATION

Supporting information is available online and contains additional SAXS results.

