## Supplementary Figures and Tables for "Structural characterization of a minimal KLC2/Nup358/BicD2 complex"

### SUPPLEMENTARY INFORMATION

#### Table S1. Summary of statistics from DAMMIF bead model 3D reconstructions of the molecular envelopes

| **Sample concentration (mg/mL)** | **Mean value of NSD** | **$\chi^2$ refined model** | **Resolution (Å)** | **# of Clusters** | **Included models (out of 15)** |
|---|---|---|---|---|---|
| **Nup358-KLC2 Fusion** | | | | | |
| 3.0 | 1.38 ±0.12 | 1.05 | 45 ±3 | 3 | 15 |
| 2.0 | 1.56 ±0.15 | 1.09 | 47 ±4 | 3 | 15 |
| 1.6 | 1.47 ±0.12 | 1.04 | 45 ±3 | 8 | 14 |
| 1.2 | 1.57 ±0.13 | 1.23 | 54 ±4 | 5 | 15 |
| 1.00 | 1.36 ±0.07 | 1.21 | 42 ±3 | 0 | 15 |
| 0.80 | 1.24 ± 0.21 | 1.08 | 40 ±3 | 4 | 14 |
| 0.60 | 1.45 ±0.11 | 1.13 | 45 ±3 | 0 | 14 |
| 0.40 | 1.54 ±0.14 | 1.08 | 51 ±4 | 4 | 14 |
| 0.20 | 0.79 ±0.20 | 1.30 | 29 ±2 | 2 | 14 |
| **Nup358-KLC2 Fusion + BicD2 in a 1:1 ratio** | | | | | |
| 3.75 | 1.52 ±0.09 | 1.09 | 58 ±4 | 7 | 14 |
| 2.50 | 1.15 ±0.09 | 1.13 | 51 ±4 | 5 | 14 |
| 2.00 | 1.06 ± 0.08 | 1.40 | 49 ±4 | 5 | 14 |
| 1.50 | 1.15 ± 0.25 | 1.14 | 54 ±4 | 9 | 14 |
| 1.25 | 1.02 ±0.09 | 1.04 | 47 ±4 | 0 | 14 |
| 1.00 | 0.76 ±0.06 | 1.09 | 41 ±3 | 7 | 14 |
| 0.75 | 0.83 ±0.13 | 1.27 | 42 ±3 | 7 | 14 |
| 0.50 | 1.62 ±0.25 | 1.45 | 54 ±4 | 8 | 14 |
| 0.25 | 1.09 ±0.21 | 1.40 | 51 ±4 | 6 | 14 |
| **Nup358/W2224A/D2225A-KLC2 Fusion** | | | | | |
| 2.00 | 1.02 ±0.13 | 1.02 | 45 ±3 | 5 | 13 |
| 1.60 | 0.86 ±0.08 | 1.11 | 39 ±3 | 4 | 14 |
| 1.20 | 0.69 ±0.06 | 1.06 | 35 ±3 | 7 | 14 |
| 1.00 | 1.06 ±0.23 | 1.03 | 41 ±3 | 5 | 14 |
| 0.80 | 0.96 ±0.12 | 1.16 | 41 ±3 | 2 | 13 |
| 0.60 | 0.82 ±0.09 | 1.12 | 37 ±3 | 4 | 14 |
| 0.40 | 0.86 ±0.11 | 1.08 | 35 ±3 | 3 | 14 |
| **Nup358/W2224A/D2225A-KLC2 Fusion+ BicD2 in a 1:1 ratio** | | | | | |
| 1.50 | 0.84 ±0.11 | 1.00 | 38 ±3 | 7 | 14 |
| 1.25 | 0.89 ±0.18 | 1.07 | 39 ±3 | 4 | 13 |
| 1.00 | 1.04 ±0.14 | 1.07 | 44 ±3 | 5 | 14 |
| 0.75 | 0.89 ±0.09 | 1.02 | 40 ±3 | 6 | 14 |
| 0.50 | 0.95 ±0.14 | 1.21 | 33 ±3 | 5 | 14 |

Where NSD: Normalized spatial discrepancy value, $\chi^2$ is the chi squared value. See also Figures S1– S30.

### Nup358-KLC2 Fusion 3 mg/mL

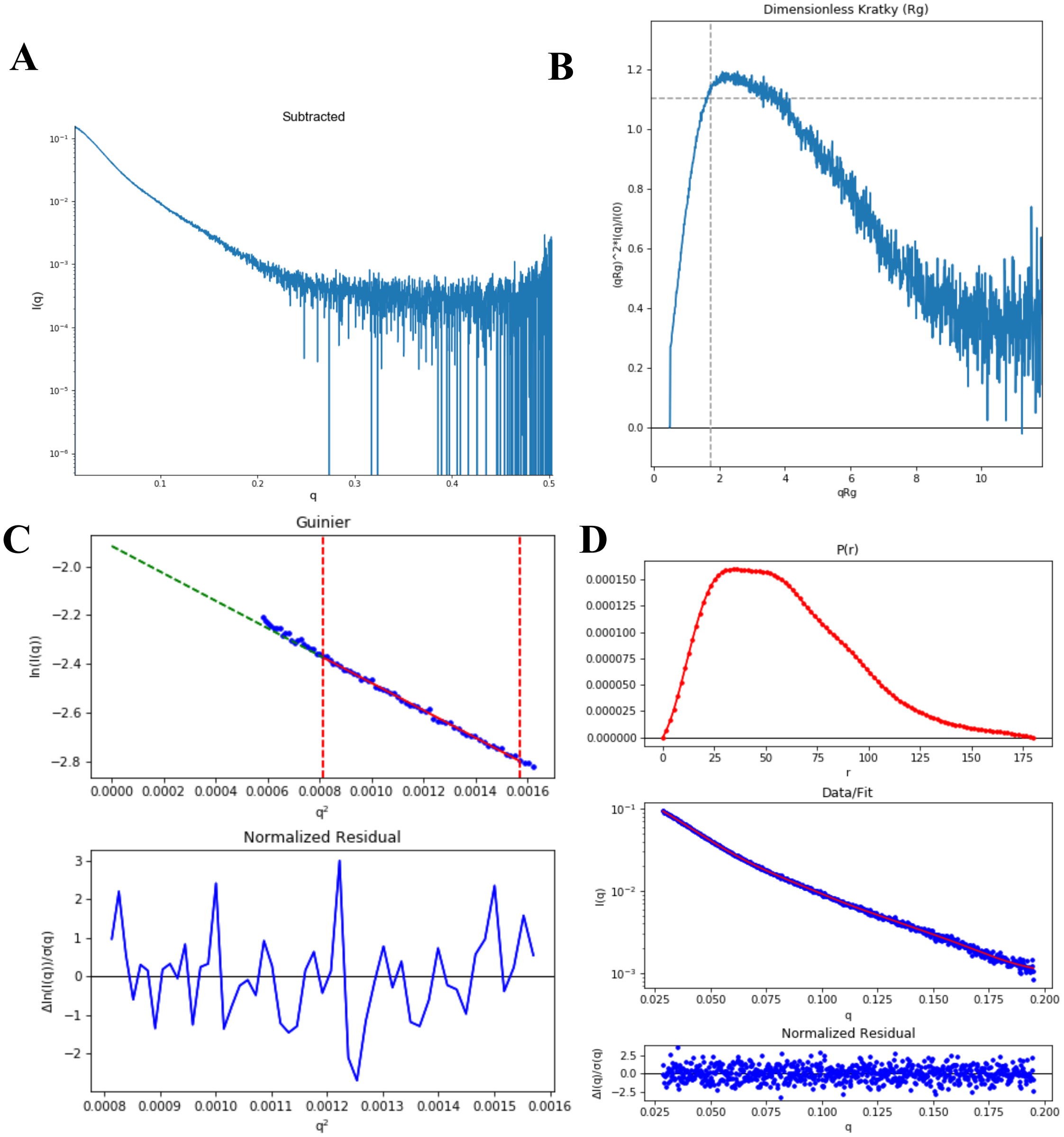


**Figure S1. SAXS plots for the Nup358-KLC2 fusion protein at 3 mg/mL.** (A) The SAXS-scattering intensity profile I (q) is shown as a function of the scattering vector (q). (B) The dimensionless Kratky plot. (C) The Guinier plot (top) is shown with the normalized residual of the Guinier fit (bottom). (D) The pair distance distribution function P(r) (top), the middle panel shows the scattering intensity profile calculated from the p(r) function (red) overlaid with the scattering intensity profile of the data (blue). The bottom panel shows the normalized residual of the fit.

**Nup358-KLC2 Fusion 2 mg/mL**

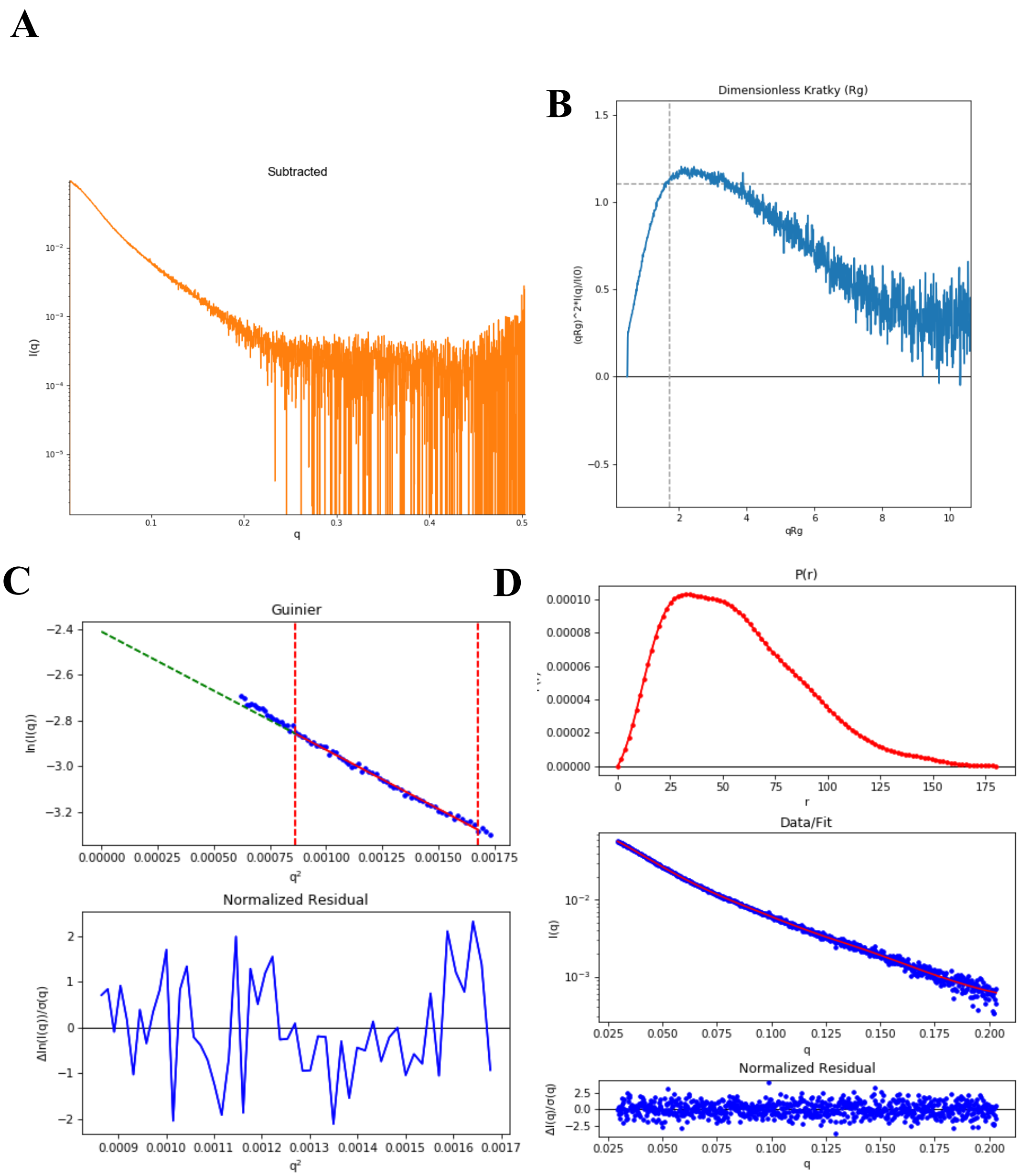


**Figure S2. SAXS plots for Nup358-KLC2 fusion 2 mg/mL.** (A) The scattering intensity profile I (q) is shown as a function of the scattering vector (q). (B) The dimensionless Kratky plot. (C) The Guinier plot (top) is shown with the normalized residual of the Guinier fit (bottom). (D) The pair distance distribution function P(r) (top), the middle panel shows the scattering intensity profile calculated from the p(r) function (red) overlaid with the scattering intensity profile of the data (blue). The bottom panel shows the normalized residual of the fit.

**Nup358-KLC2 Fusion 1.6 mg/mL**

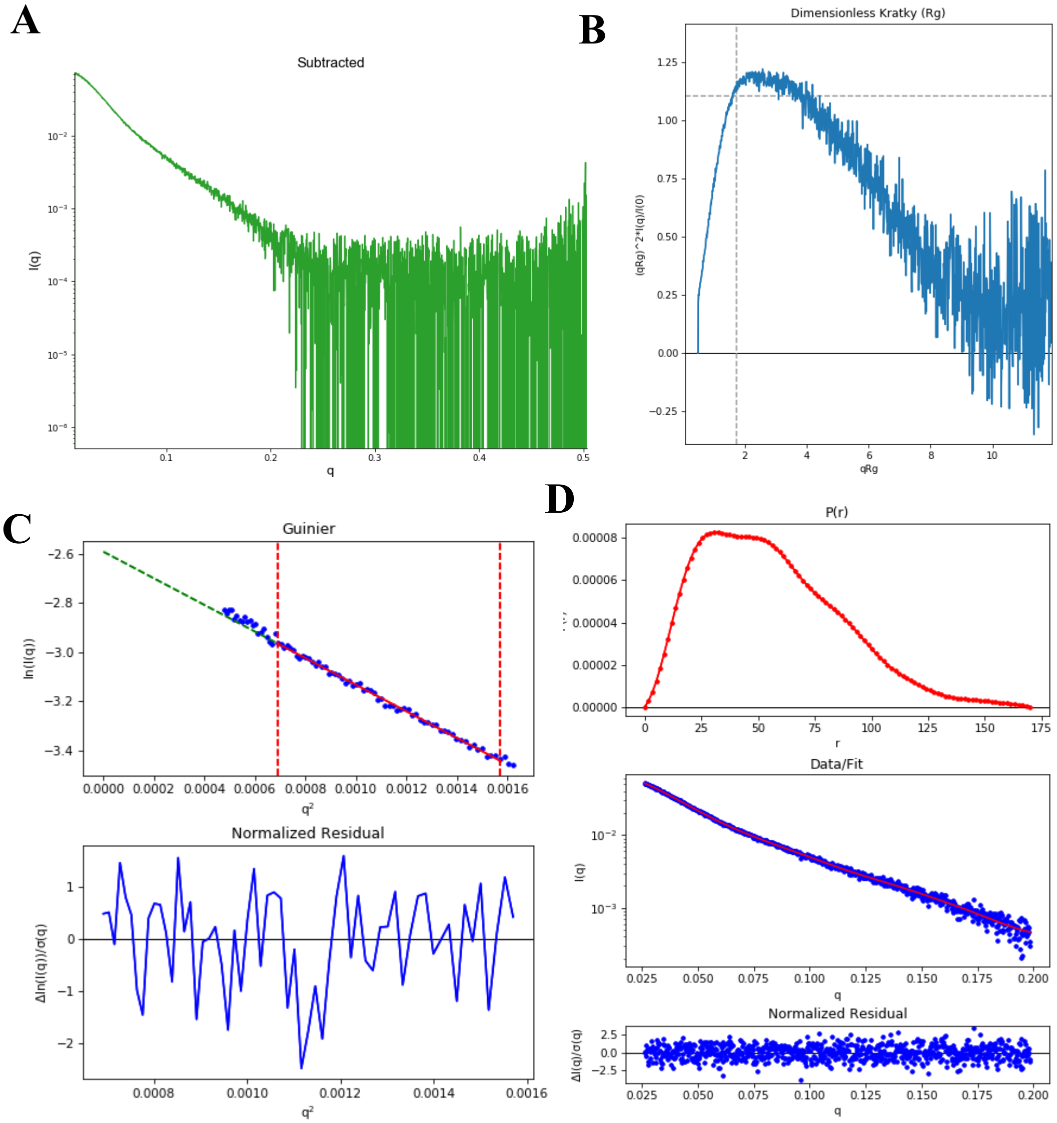


**Figure S3. SAXS plots for Nup358-KLC2 fusion 1.6 mg/mL.** (A) The scattering intensity profile I (q) is shown as a function of the scattering vector (q). (B) The dimensionless Kratky plot. (C) The Guinier plot (top) is shown with the normalized residual of the Guinier fit (bottom). (D) The pair distance distribution function P(r) (top), the middle panel shows the scattering intensity profile calculated from the p(r) function (red) overlaid with the scattering intensity profile of the data (blue). The bottom panel shows the normalized residual of the fit.

### Nup358-KLC2 Fusion 1.2 mg/mL

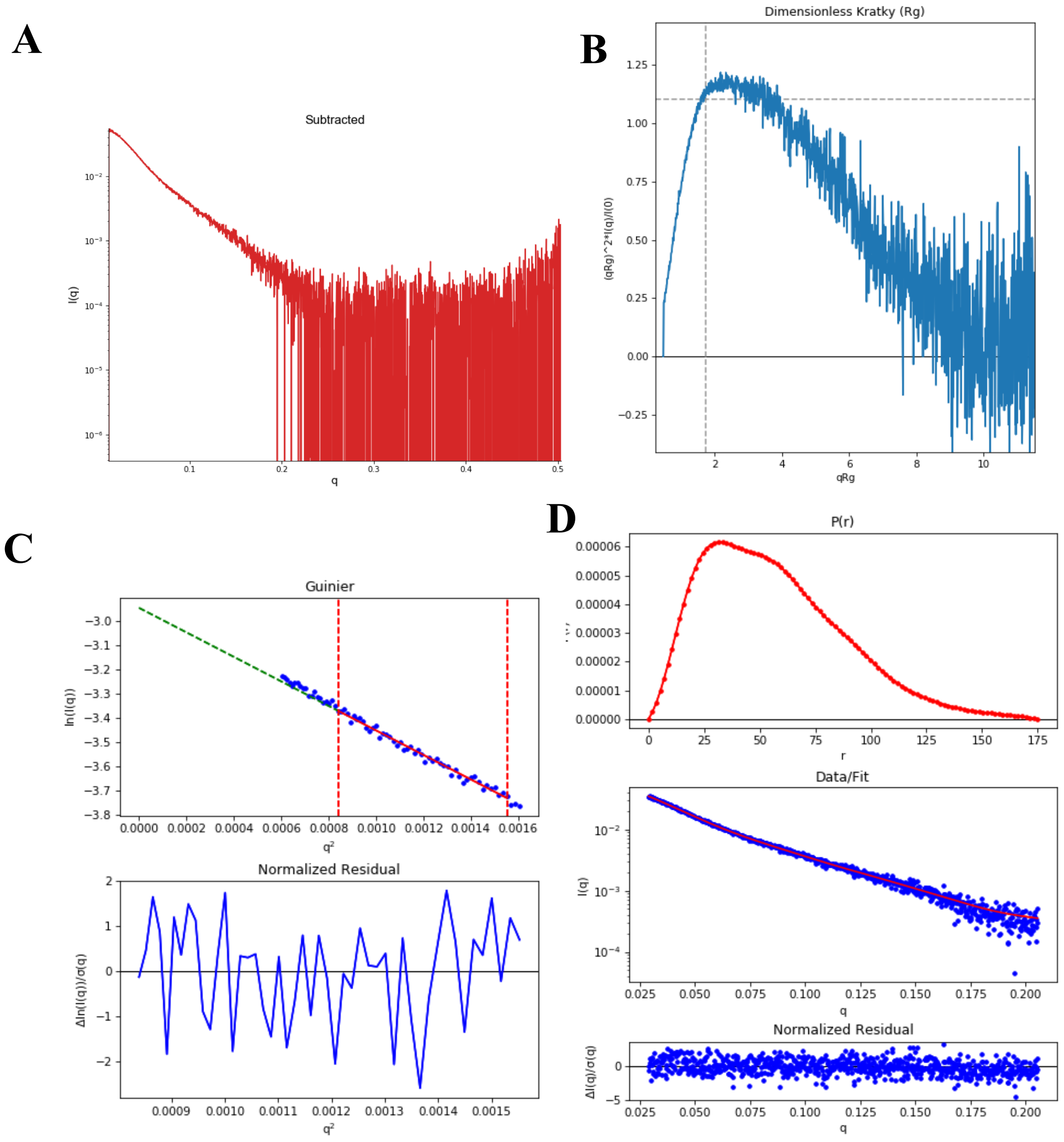


**Figure S4. SAXS plots for Nup358-KLC2 fusion 1.2 mg/mL.** (A) The scattering intensity profile I (q) is shown as a function of the scattering vector (q). (B) The dimensionless Kratky plot. (C) The Guinier plot (top) is shown with the normalized residual of the Guinier fit (bottom). (D) The pair distance distribution function P(r) (top), the middle panel shows the scattering intensity profile calculated from the p(r) function (red) overlaid with the scattering intensity profile of the data (blue). The bottom panel shows the normalized residual of the fit.

### Nup358-KLC2 Fusion 1 mg/mL

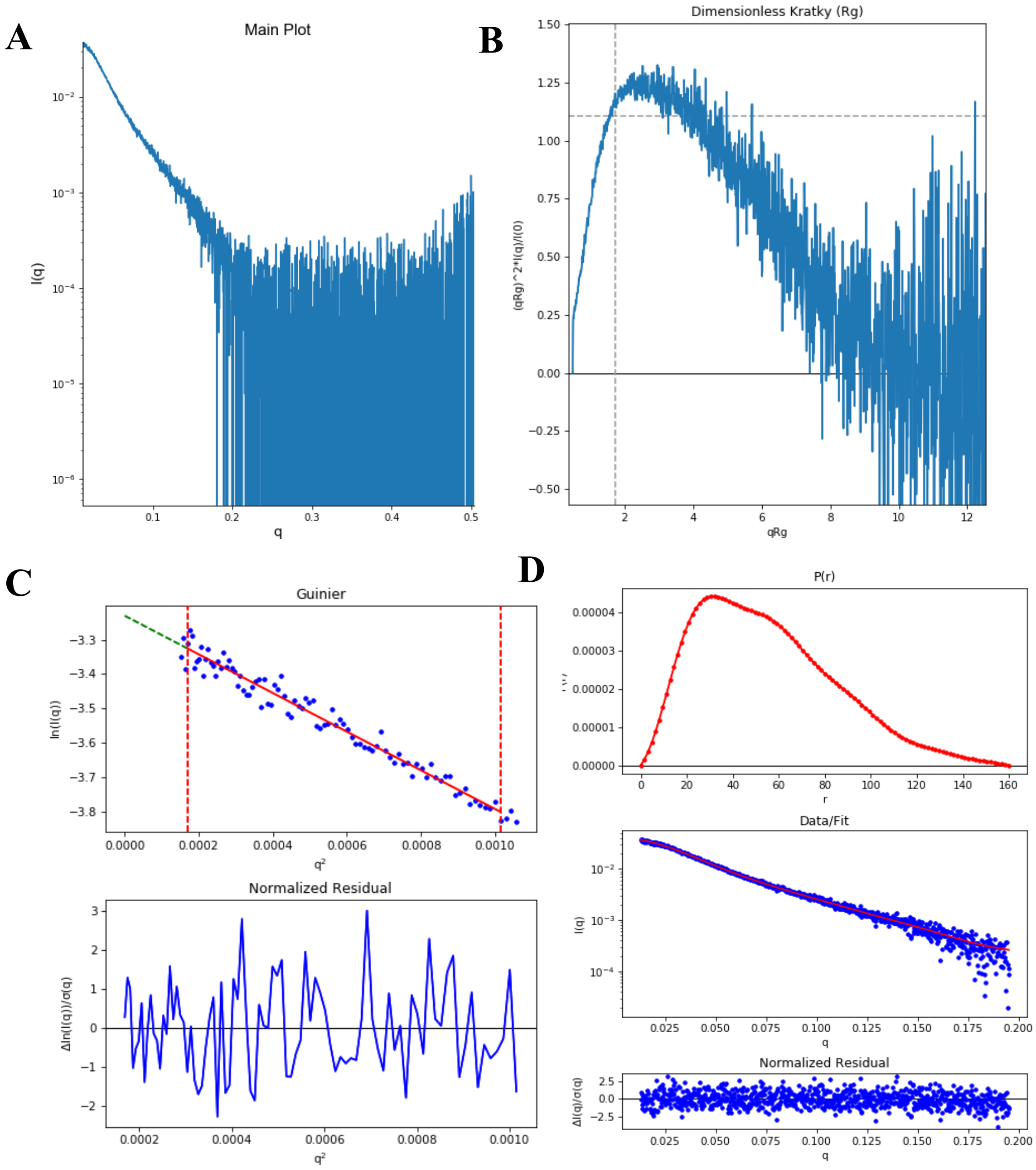


**Figure S5. SAXS plots for Nup358-KLC2 fusion 1 mg/mL.** (A) The scattering intensity profile I (q) is shown as a function of the scattering vector (q). (B) The dimensionless Kratky plot. (C) The Guinier plot (top) is shown with the normalized residual of the Guinier fit (bottom). (D) The pair distance distribution function P(r) (top), the middle panel shows the scattering intensity profile calculated from the p(r) function (red) overlaid with the scattering intensity profile of the data (blue). The bottom panel shows the normalized residual of the fit.

### Nup358-KLC2 Fusion 0.8 mg/mL

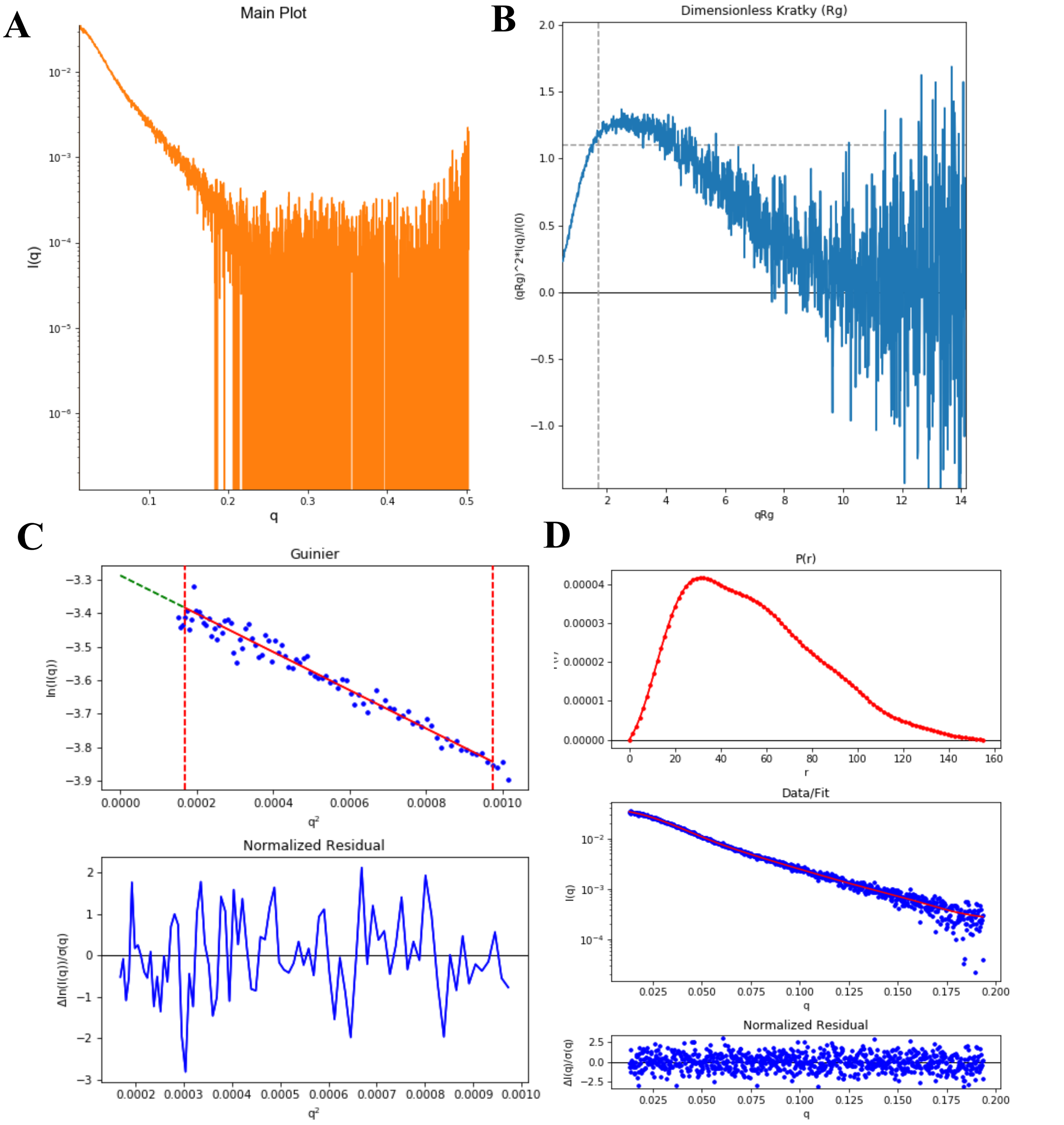


**Figure S6. SAXS plots for Nup358-KLC2 fusion 0.8 mg/mL.** (A) The scattering intensity profile I (q) is shown as a function of the scattering vector (q). (B) The dimensionless Kratky plot. (C) The Guinier plot (top) is shown with the normalized residual of the Guinier fit (bottom). (D) The pair distance distribution function P(r) (top), the middle panel shows the scattering intensity profile calculated from the p(r) function (red) overlaid with the scattering intensity profile of the data (blue). The bottom panel shows the normalized residual of the fit.

**Nup358-KLC2 Fusion 0.6 mg/mL**

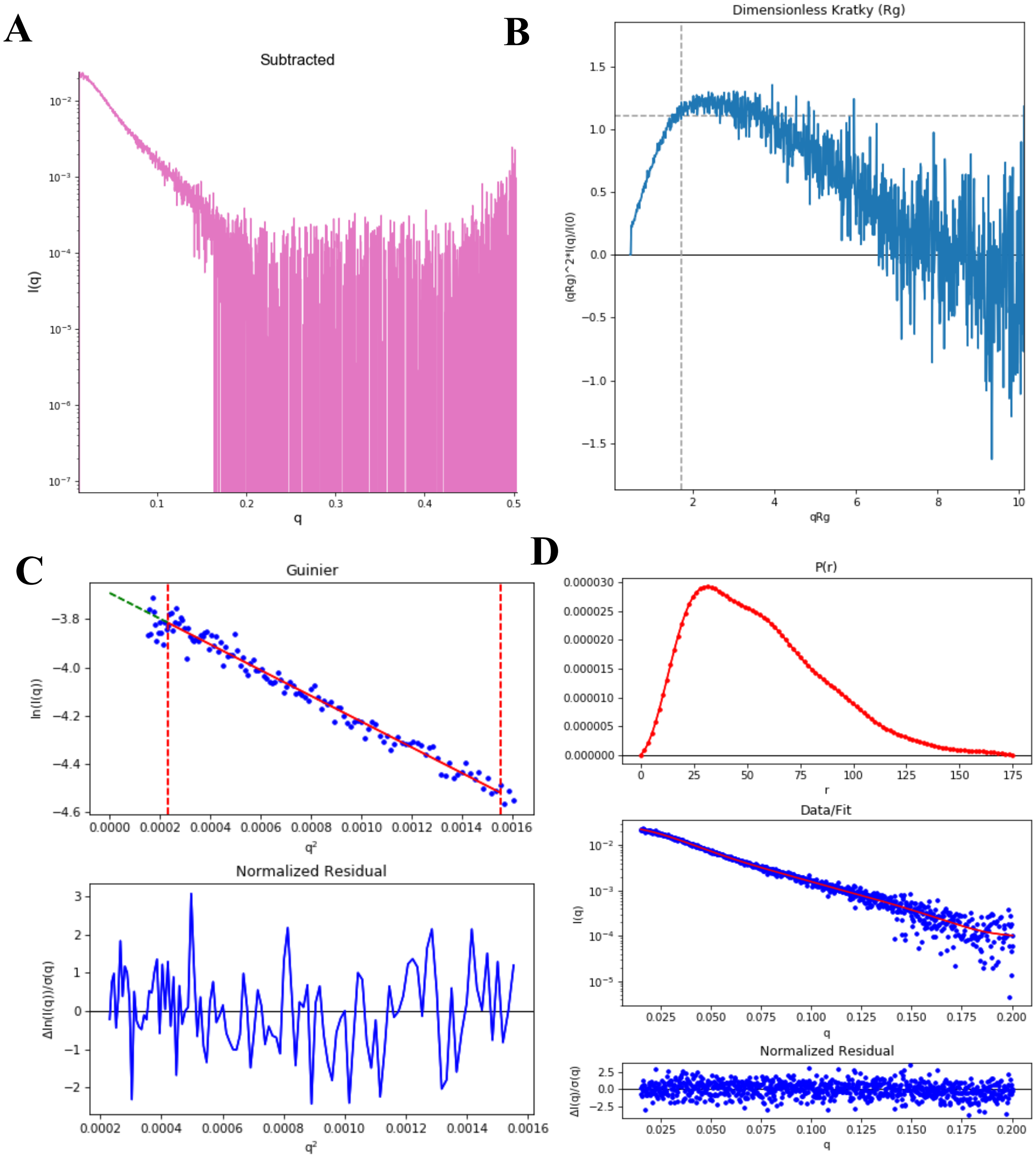


**Figure S7. SAXS plots for Nup358-KLC2 fusion 0.6 mg/mL.** (A) The scattering intensity profile I (q) is shown as a function of the scattering vector (q). (B) The dimensionless Kratky plot. (C) The Guinier plot (top) is shown with the normalized residual of the Guinier fit (bottom). (D) The pair distance distribution function P(r) (top), the middle panel shows the scattering intensity profile calculated from the p(r) function (red) overlaid with the scattering intensity profile of the data (blue). The bottom panel shows the normalized residual of the fit.

### Nup358-KLC2 Fusion 0.4 mg/mL

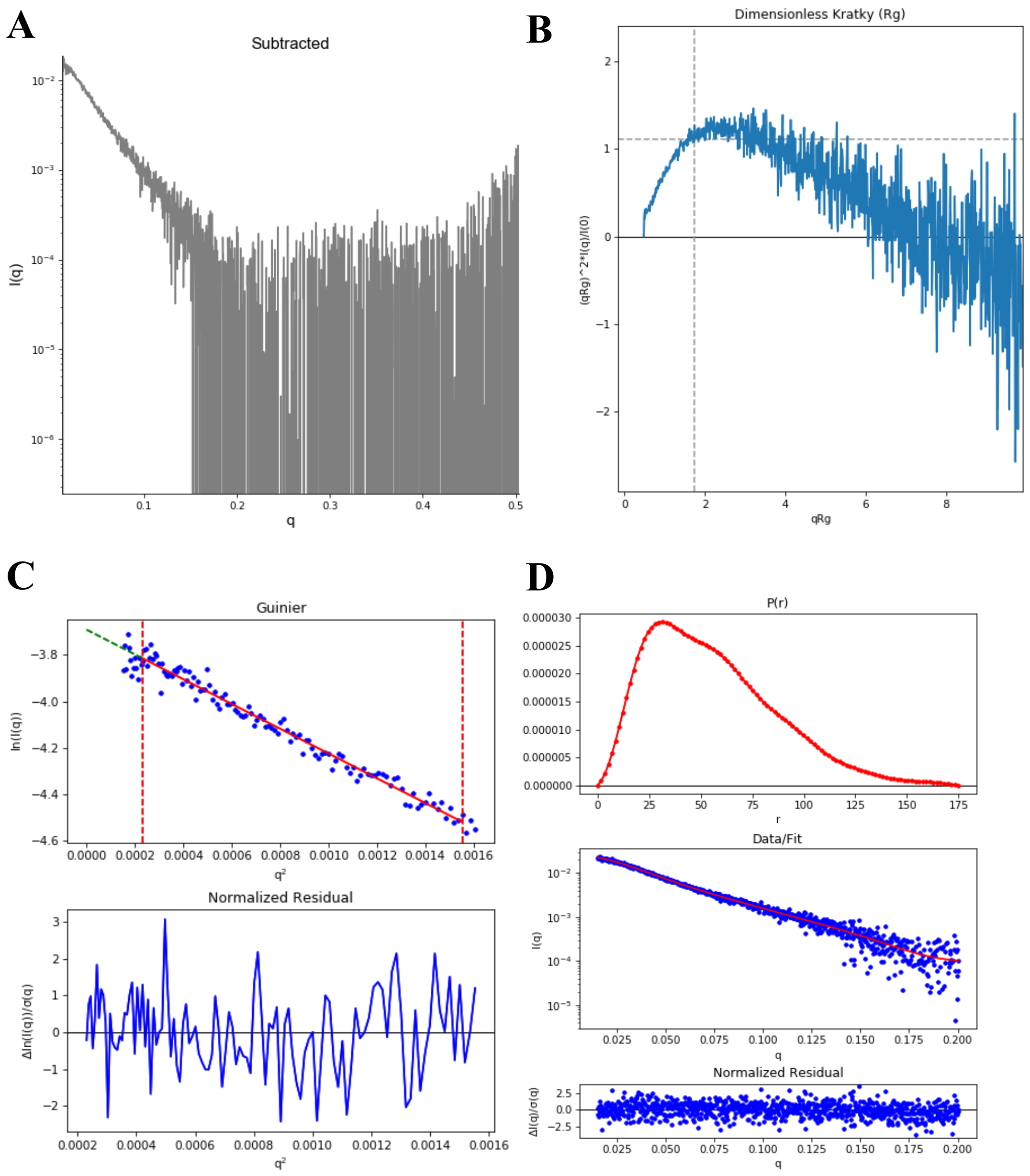


**Figure S8. SAXS plots for Nup358-KLC2 fusion 0.4 mg/mL.** (A) The scattering intensity profile I (q) is shown as a function of the scattering vector (q). (B) The dimensionless Kratky plot. (C) The Guinier plot (top) is shown with the normalized residual of the Guinier fit (bottom). (D) The pair distance distribution function P(r) (top), the middle panel shows the scattering intensity profile calculated from the p(r) function (red) overlaid with the scattering intensity profile of the data (blue). The bottom panel shows the normalized residual of the fit.

**Nup358-KLC2 fusion 0.2 mg/mL**

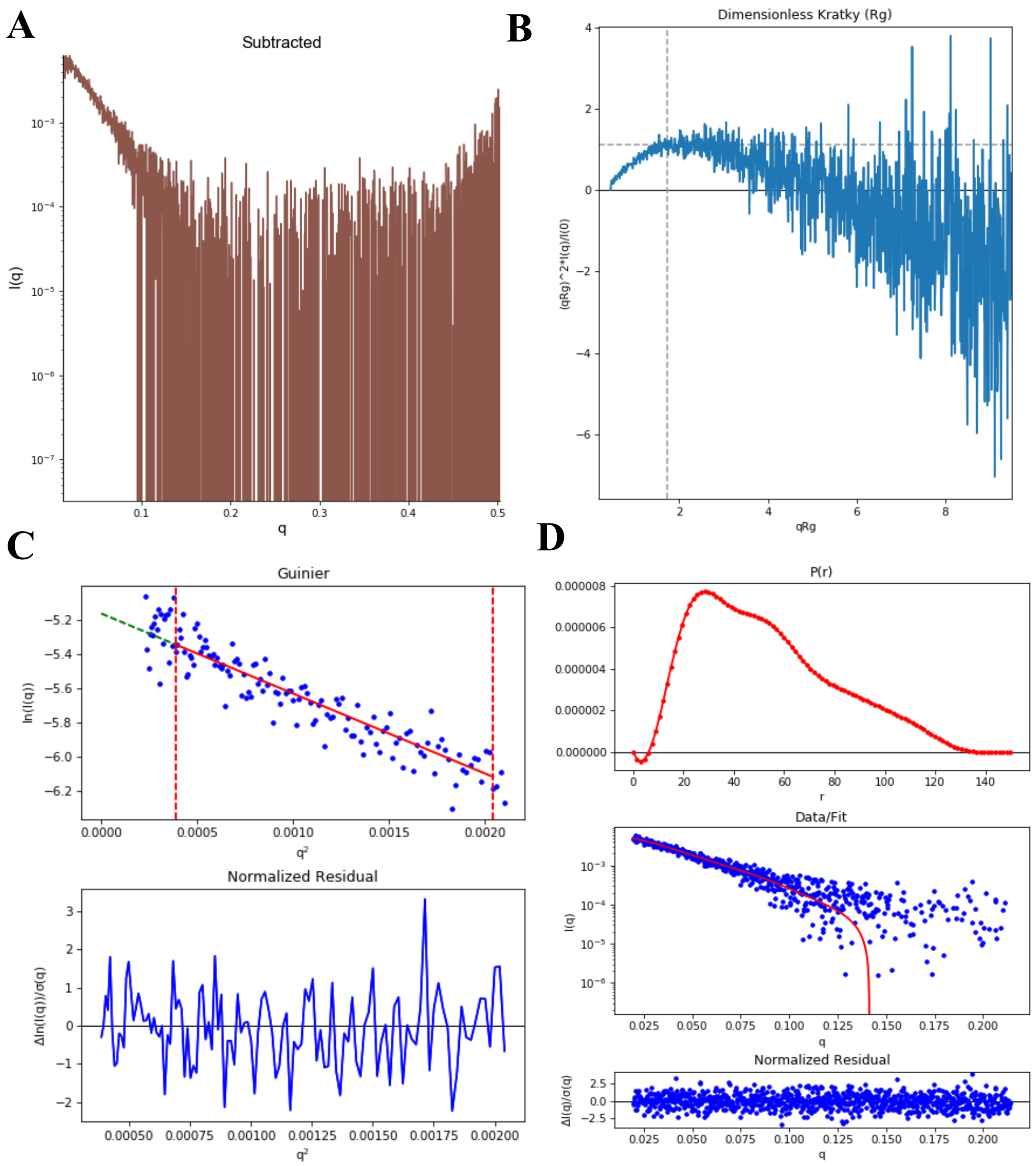


**Figure S9. SAXS plots for Nup358-KLC2 fusion 0.2 mg/mL.** (A) The scattering intensity profile I (q) is shown as a function of the scattering vector (q). (B) The dimensionless Kratky plot. (C) The Guinier plot (top) is shown with the normalized residual of the Guinier fit (bottom). (D) The pair distance distribution function P(r) (top), the middle panel shows the scattering intensity profile calculated from the p(r) function (red) overlaid with the scattering intensity profile of the data (blue). The bottom panel shows the normalized residual of the fit.

**Nup358-KLC2 Fusion 3 mg/mL + BicD2 0.75mg/mL**

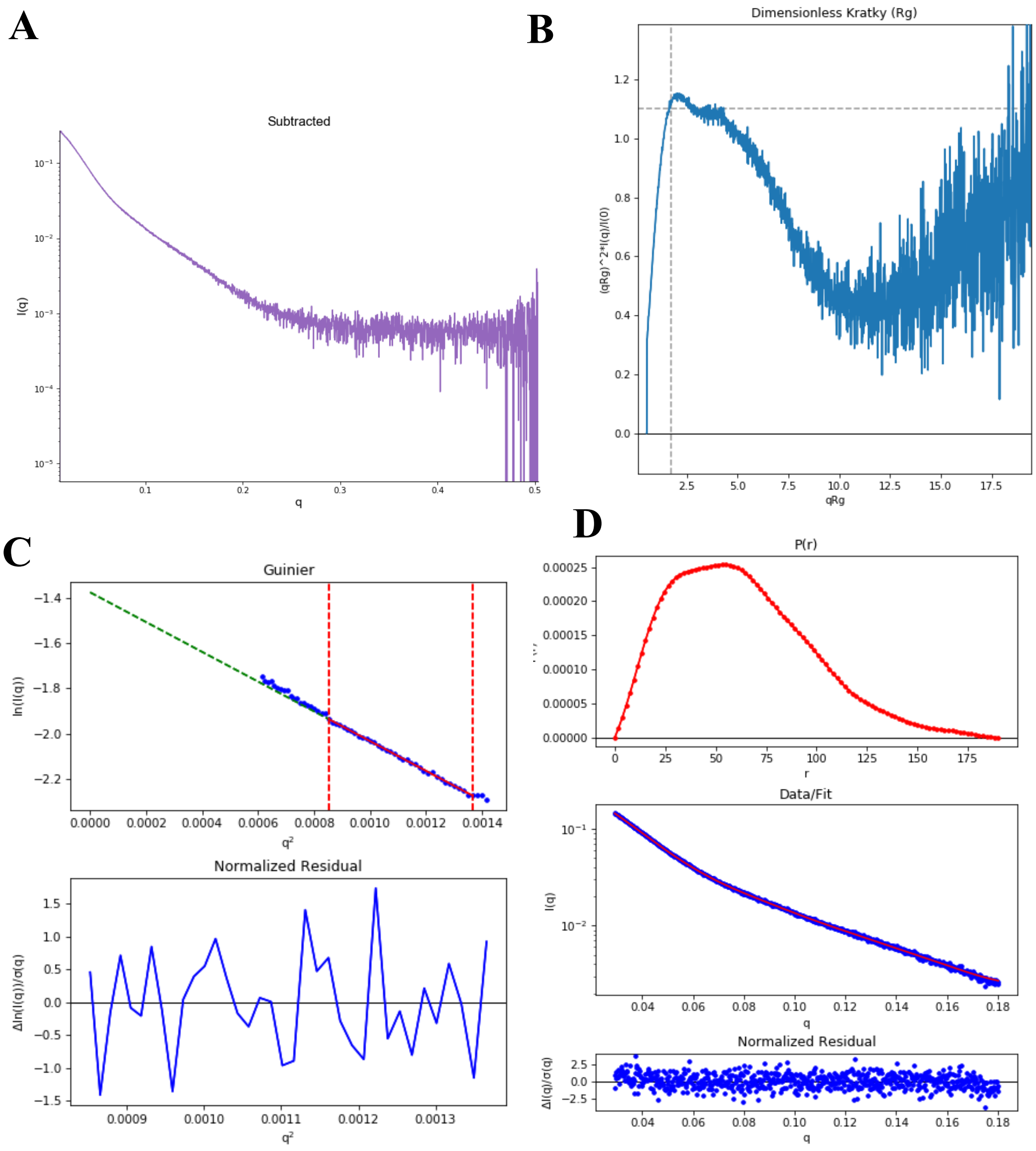


**Figure S10. SAXS plots for Nup358-KLC2 Fusion 3 mg/mL + BicD2 0.75 mg/mL.** (A) The scattering intensity profile I (q) is shown as a function of the scattering vector (q). (B) The dimensionless Kratky plot. (C) The Guinier plot (top) is shown with the normalized residual of the Guinier fit (bottom). (D) The pair distance distribution function P(r) (top), the middle panel shows the scattering intensity profile calculated from the p(r) function (red) overlaid with the scattering intensity profile of the data (blue). The bottom panel shows the normalized residual of the fit.

### Nup358-KLC2 Fusion 2 mg/mL + BicD2-CTD 0.5 mg/mL

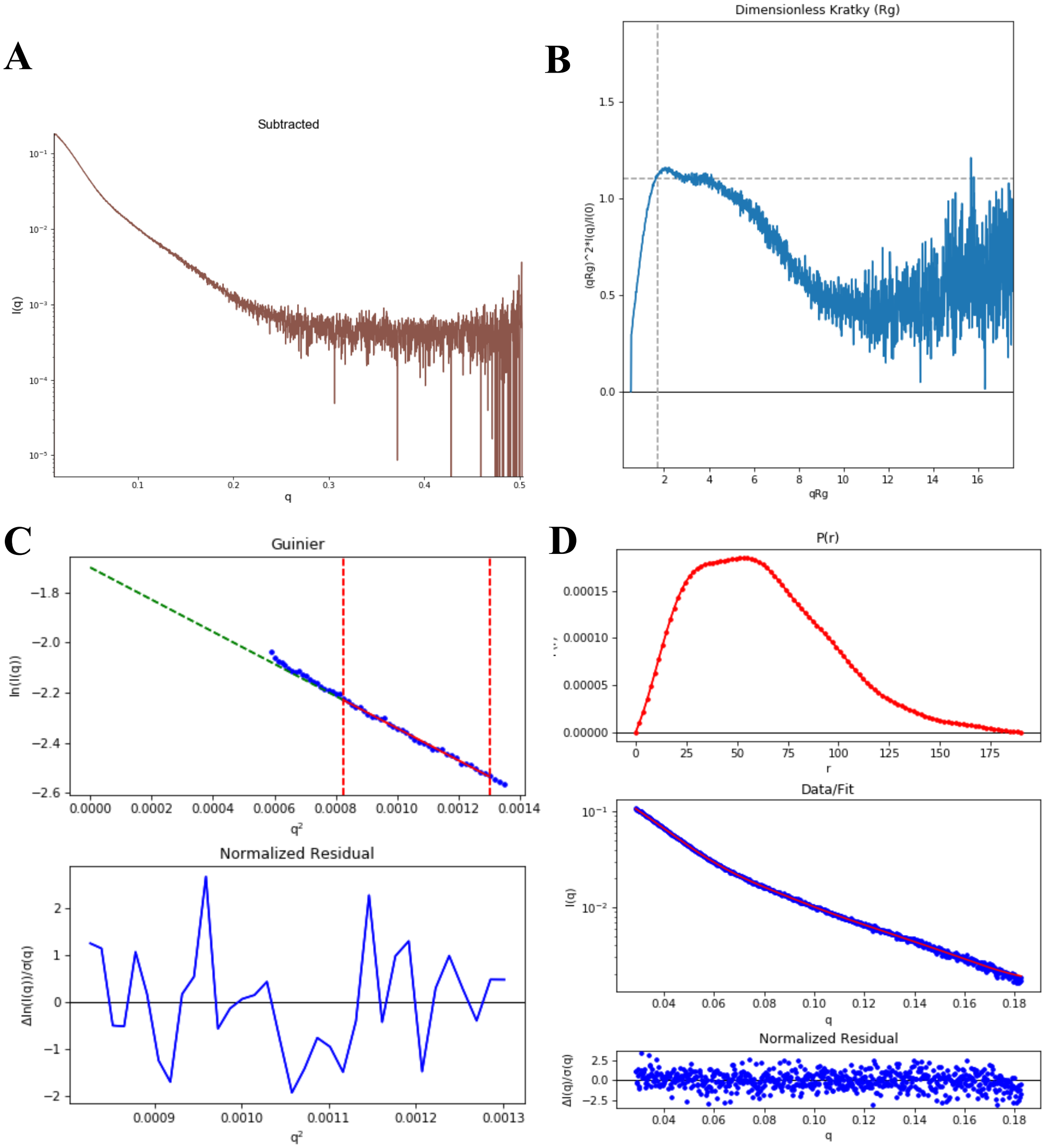


**Figure S11. SAXS plots for Nup358-KLC2 Fusion 2 mg/mL + BicD2-CTD 0.5 mg/mL.** (A) The scattering intensity profile I (q) is shown as a function of the scattering vector (q). (B) The dimensionless Kratky plot. (C) The Guinier plot (top) is shown with the normalized residual of the Guinier fit (bottom). (D) The pair distance distribution function P(r) (top), the middle panel shows the scattering intensity profile calculated from the p(r) function (red) overlaid with the scattering intensity profile of the data (blue). The bottom panel shows the normalized residual of the fit.

**Nup358-KLC2 Fusion 1.6 mg/mL + BicD2-CTD 0.4 mg/mL**

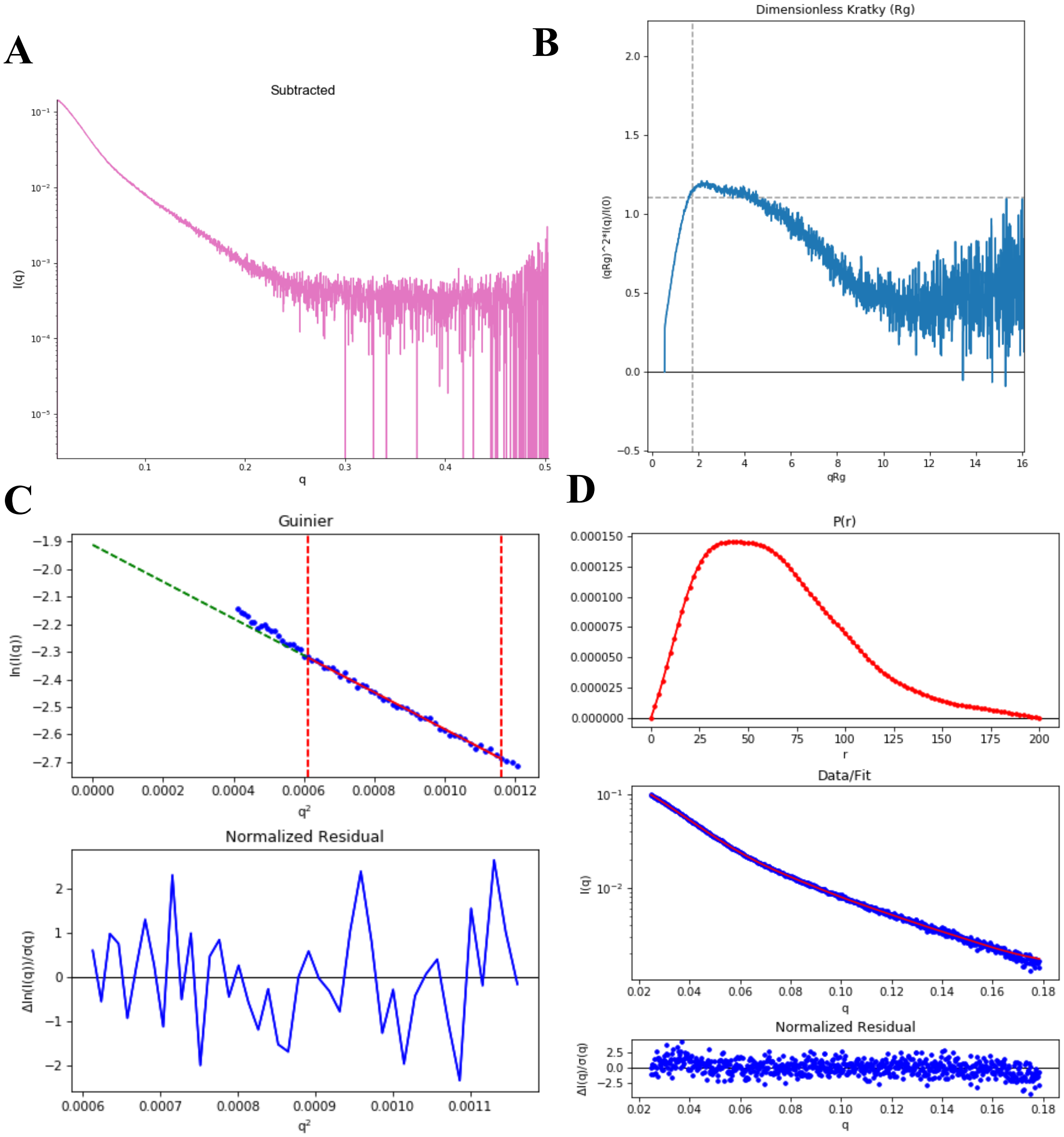


**Figure S12. SAXS plots for Nup358-KLC2 Fusion 1.6 mg/mL + BicD2-CTD 0.4 mg/mL.** (A) The scattering intensity profile I (q) is shown as a function of the scattering vector (q). (B) The dimensionless Kratky plot. (C) The Guinier plot (top) is shown with the normalized residual of the Guinier fit (bottom). (D) The pair distance distribution function P(r) (top), the middle panel shows the scattering intensity profile calculated from the p(r) function (red) overlaid with the scattering intensity profile of the data (blue). The bottom panel shows the normalized residual of the fit.

**Nup358-KLC2 Fusion 1.2 mg/mL + BicD2-CTD 0.3 mg/mL**

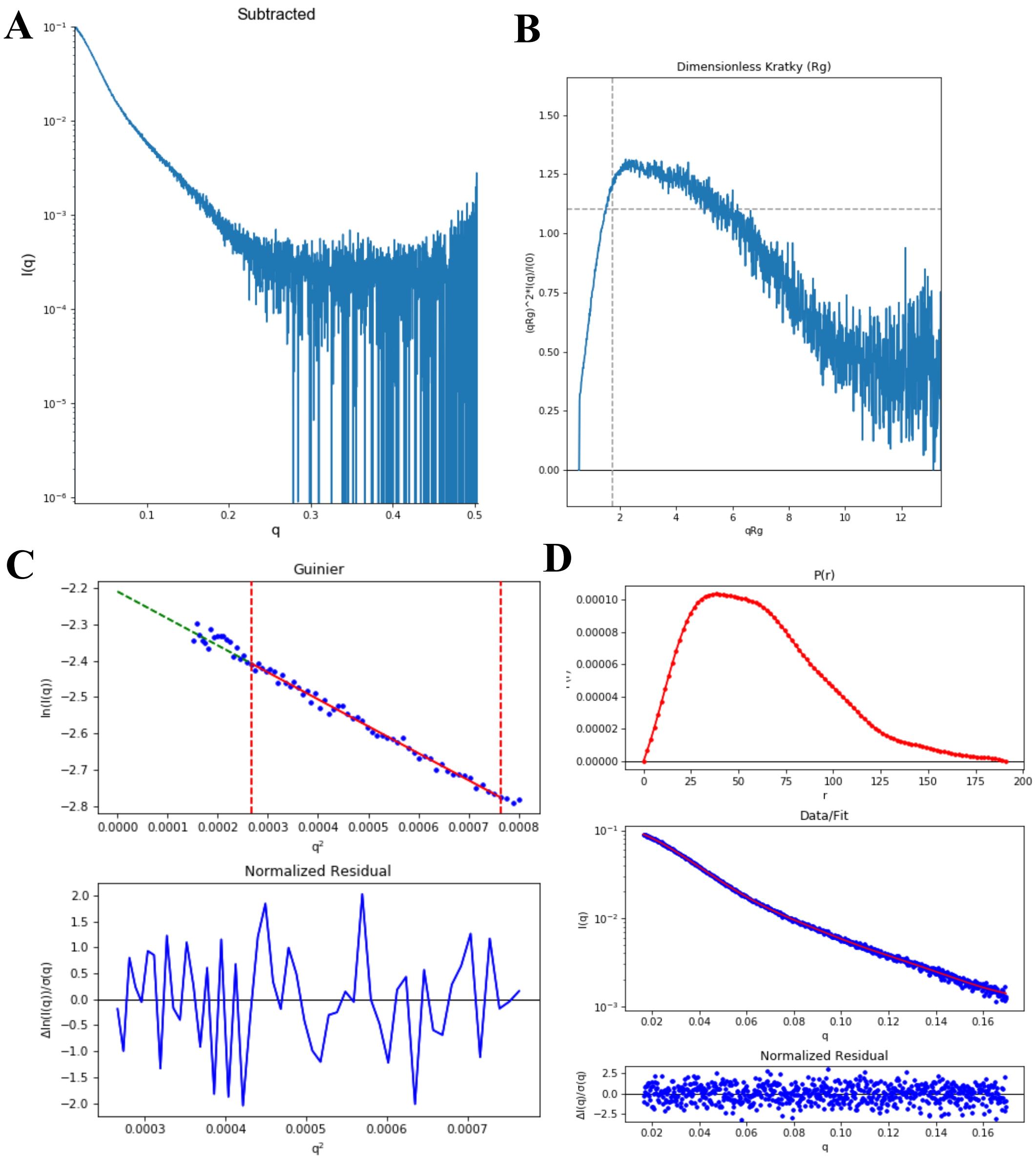


**Figure S13. SAXS plots for Nup358-KLC2 Fusion 1.2 mg/mL + BicD2-CTD 0.3 mg/mL.** (A) The scattering intensity profile I (q) is shown as a function of the scattering vector (q). (B) The dimensionless Kratky plot. (C) The Guinier plot (top) is shown with the normalized residual of the Guinier fit (bottom). (D) The pair distance distribution function P(r) (top), the middle panel shows the scattering intensity profile calculated from the p(r) function (red) overlaid with the scattering intensity profile of the data (blue). The bottom panel shows the normalized residual of the fit.

**Nup358-KLC2 Fusion 0.8 mg/mL + BicD2-CTD 0.2 mg/mL**

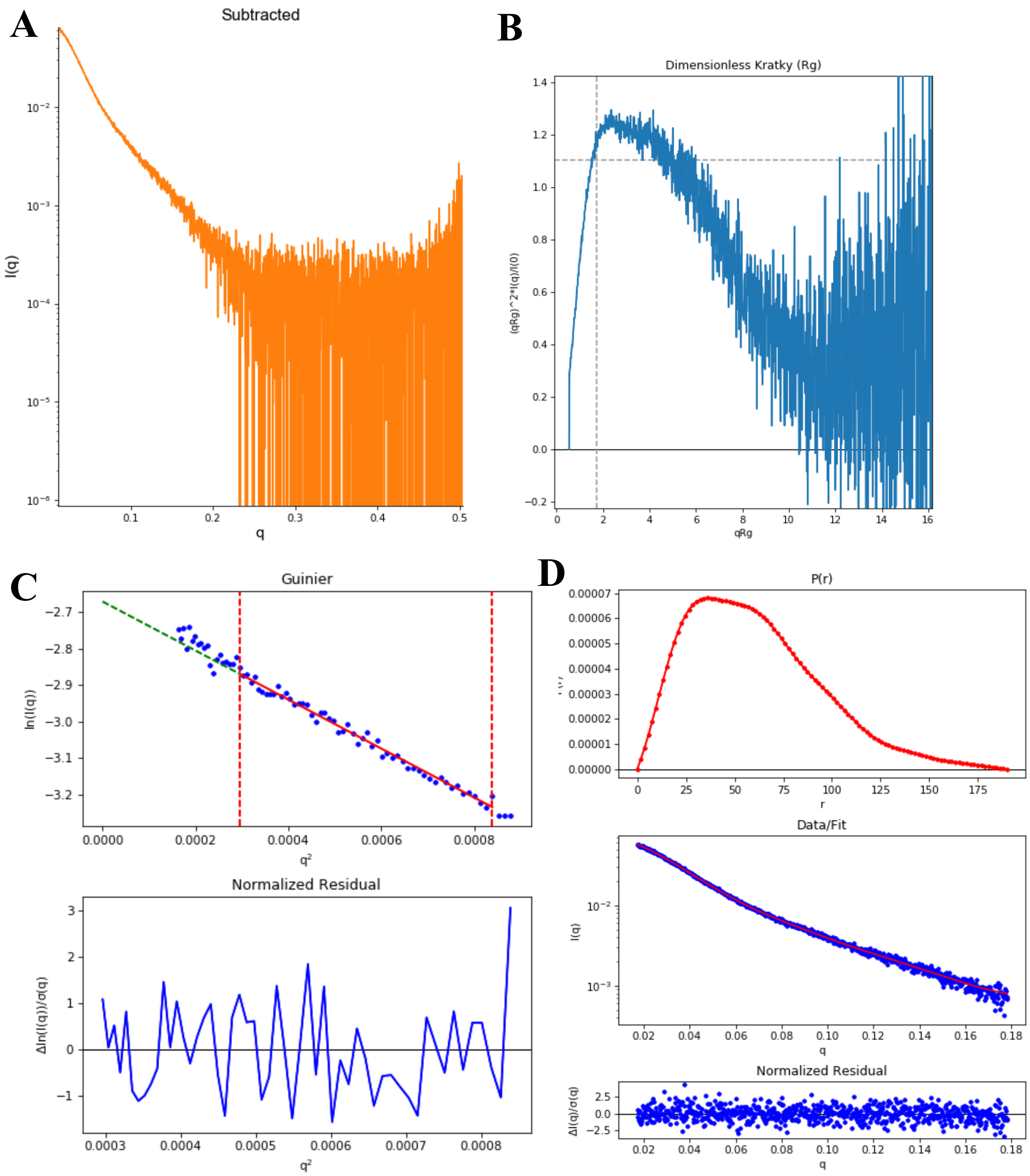


**Figure S14. SAXS plots for Nup358-KLC2 Fusion 0.8 mg/mL + BicD2-CTD 0.2 mg/mL.** (A) The scattering intensity profile I (q) is shown as a function of the scattering vector (q). (B) The dimensionless Kratky plot. (C) The Guinier plot (top) is shown with the normalized residual of the Guinier fit (bottom). (D) The pair distance distribution function P(r) (top), the middle panel shows the scattering intensity profile calculated from the p(r) function (red) overlaid with the scattering intensity profile of the data (blue). The bottom panel shows the normalized residual of the fit.

**Nup358-KLC2 fusion 0.6 mg/mL + BicD2-CTD 0.125 mg/mL**

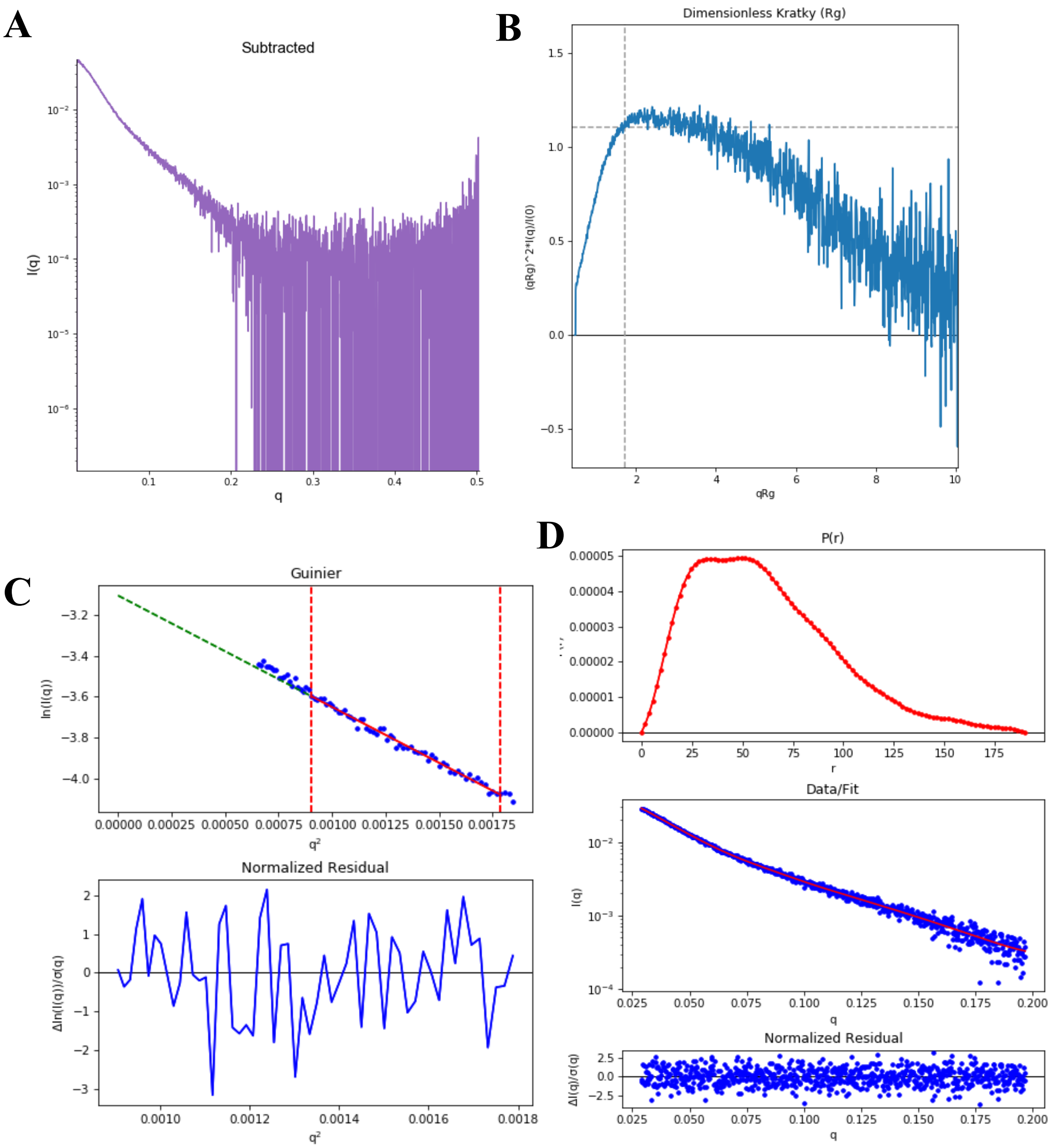


**Figure S15. SAXS plots for Nup358-KLC2 Fusion 0.6 mg/mL + BicD2-CTD 0.125 mg/mL.** (A) The scattering intensity profile I (q) is shown as a function of the scattering vector (q). (B) The dimensionless Kratky plot. (C) The Guinier plot (top) is shown with the normalized residual of the Guinier fit (bottom). (D) The pair distance distribution function P(r) (top), the middle panel shows the scattering intensity profile calculated from the p(r) function (red) overlaid with the scattering intensity profile of the data (blue). The bottom panel shows the normalized residual of the fit.

**Nup358-KLC2 fusion 0.4 mg/mL + BicD2-CTD 0.1 mg/mL**

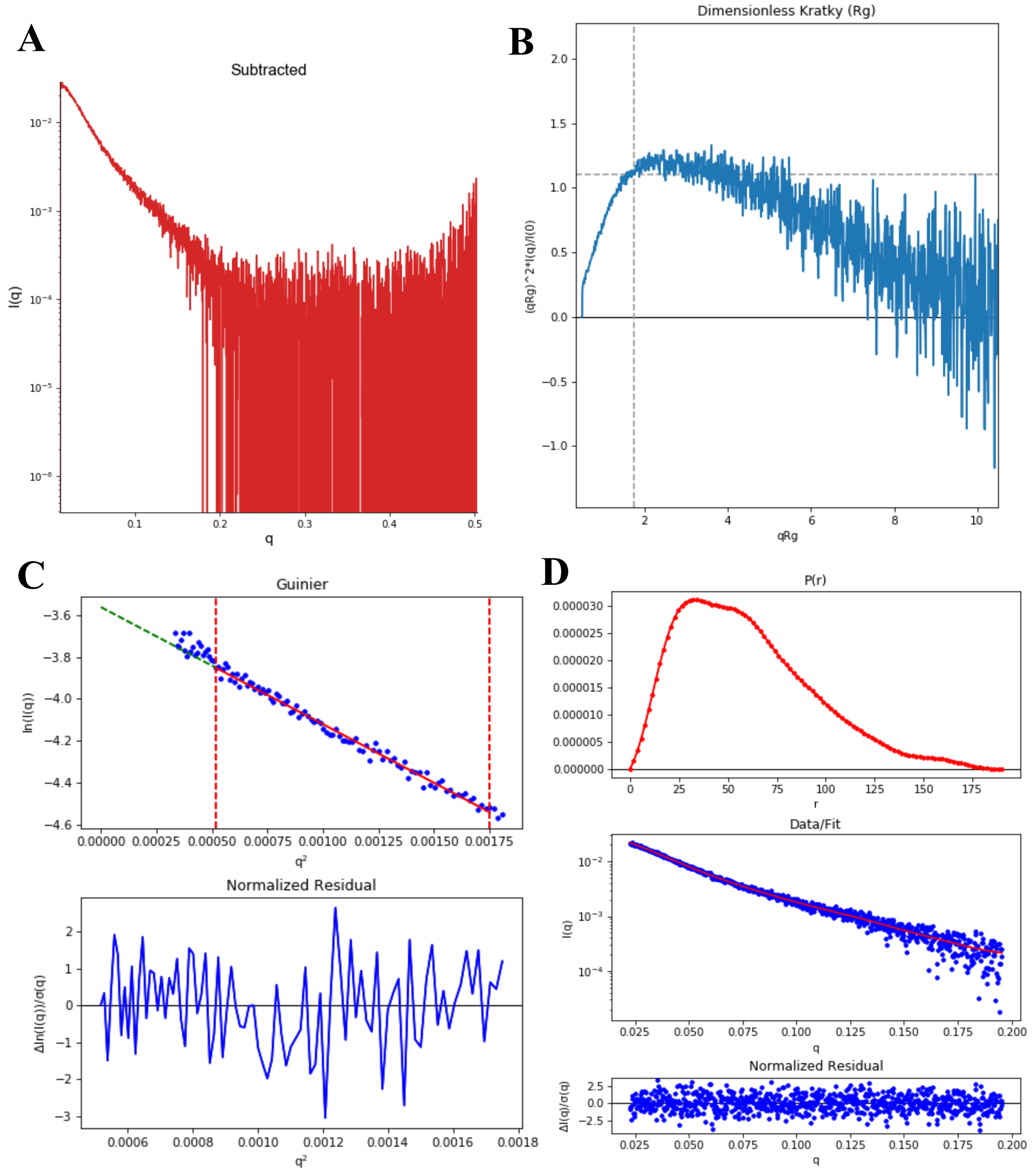


**Figure S16. SAXS plots for Nup358-KLC2 Fusion 0.4 mg/mL + BicD2-CTD 0.1 mg/mL.** (A) The scattering intensity profile I (q) is shown as a function of the scattering vector (q). (B) The dimensionless Kratky plot. (C) The Guinier plot (top) is shown with the normalized residual of the Guinier fit (bottom). (D) The pair distance distribution function P(r) (top), the middle panel shows the scattering intensity profile calculated from the p(r) function (red) overlaid with the scattering intensity profile of the data (blue). The bottom panel shows the normalized residual of the fit.

**Nup358-KLC2 fusion 0.2 mg/mL + BicD2-CTD 0.05 mg/mL**

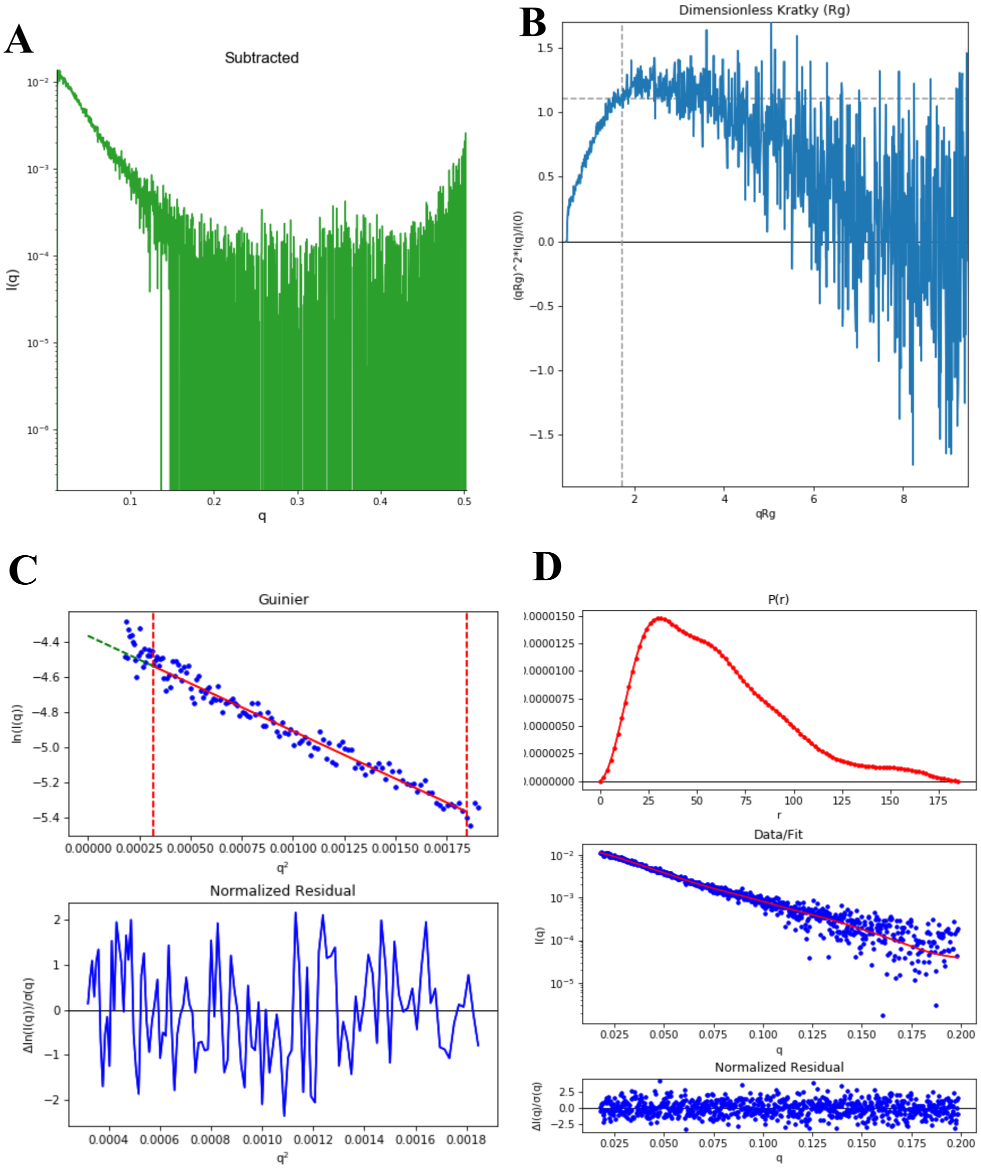


**Figure S17. SAXS plots for Nup358-KLC2 Fusion 0.2 mg/mL + BicD2-CTD 0.05 mg/mL.** (A) The scattering intensity profile I (q) is shown as a function of the scattering vector (q). (B) The dimensionless Kratky plot. (C) The Guinier plot (top) is shown with the normalized residual of the Guinier fit (bottom). (D) The pair distance distribution function P(r) (top), the middle panel shows the scattering intensity profile calculated from the p(r) function (red) overlaid with the scattering intensity profile of the data (blue). The bottom panel shows the normalized residual of the fit.

**Nup358-KLC2/W2224A/D2225A fusion 2 mg/mL**

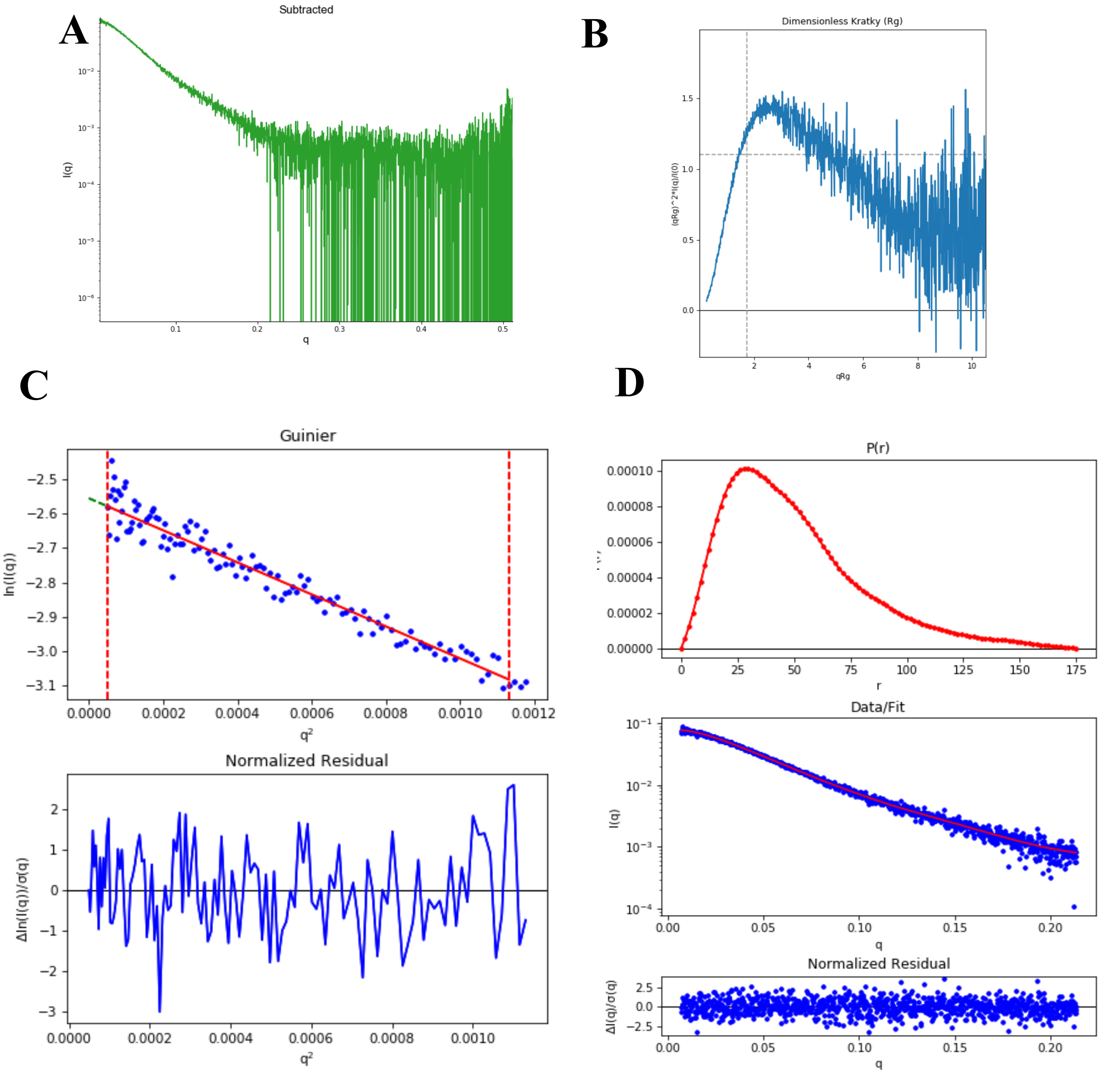


**Figure S18. SAXS plots for Nup358-KLC2/W2224A/D2225A 2 mg/mL.** (A) The scattering intensity profile I (q) is shown as a function of the scattering vector (q). (B) The dimensionless Kratky plot. (C) The Guinier plot (top) is shown with the normalized residual of the Guinier fit (bottom). (D) The pair distance distribution function P(r) (top), the middle panel shows the scattering intensity profile calculated from the p(r) function (red) overlaid with the scattering intensity profile of the data (blue). The bottom panel shows the normalized residual of the fit.

**Nup358-KLC2/W2224A/D2225A fusion 1.6 mg/mL**

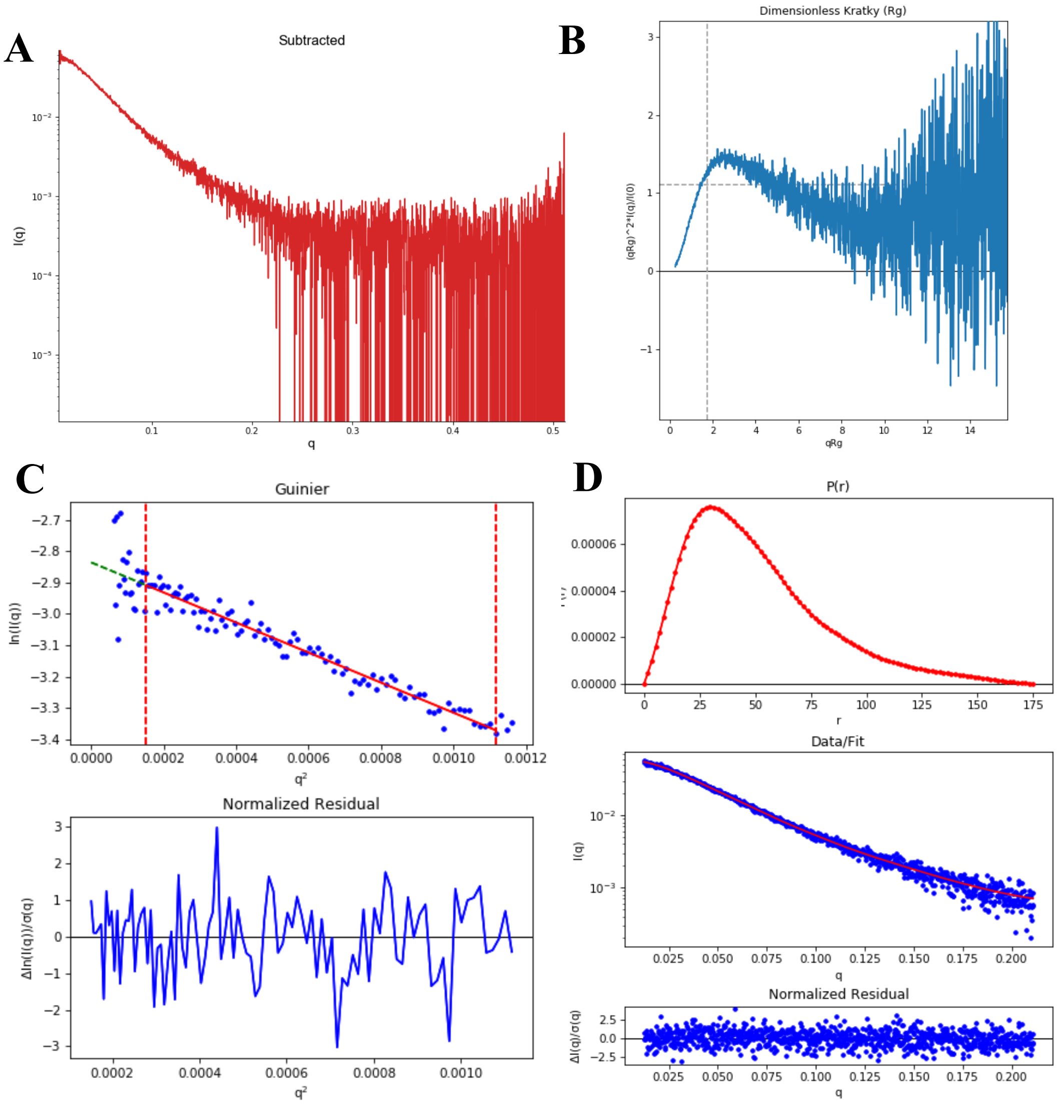


**Figure S19. SAXS plots for Nup358-KLC2/W2224A/D2225A 1.6 mg/mL.** (A) The scattering intensity profile I (q) is shown as a function of the scattering vector (q). (B) The dimensionless Kratky plot. (C) The Guinier plot (top) is shown with the normalized residual of the Guinier fit (bottom). (D) The pair distance distribution function P(r) (top), the middle panel shows the scattering intensity profile calculated from the p(r) function (red) overlaid with the scattering intensity profile of the data (blue). The bottom panel shows the normalized residual of the fit.

**Nup358-KLC2/W2224A/D2225A fusion 1.2 mg/mL**

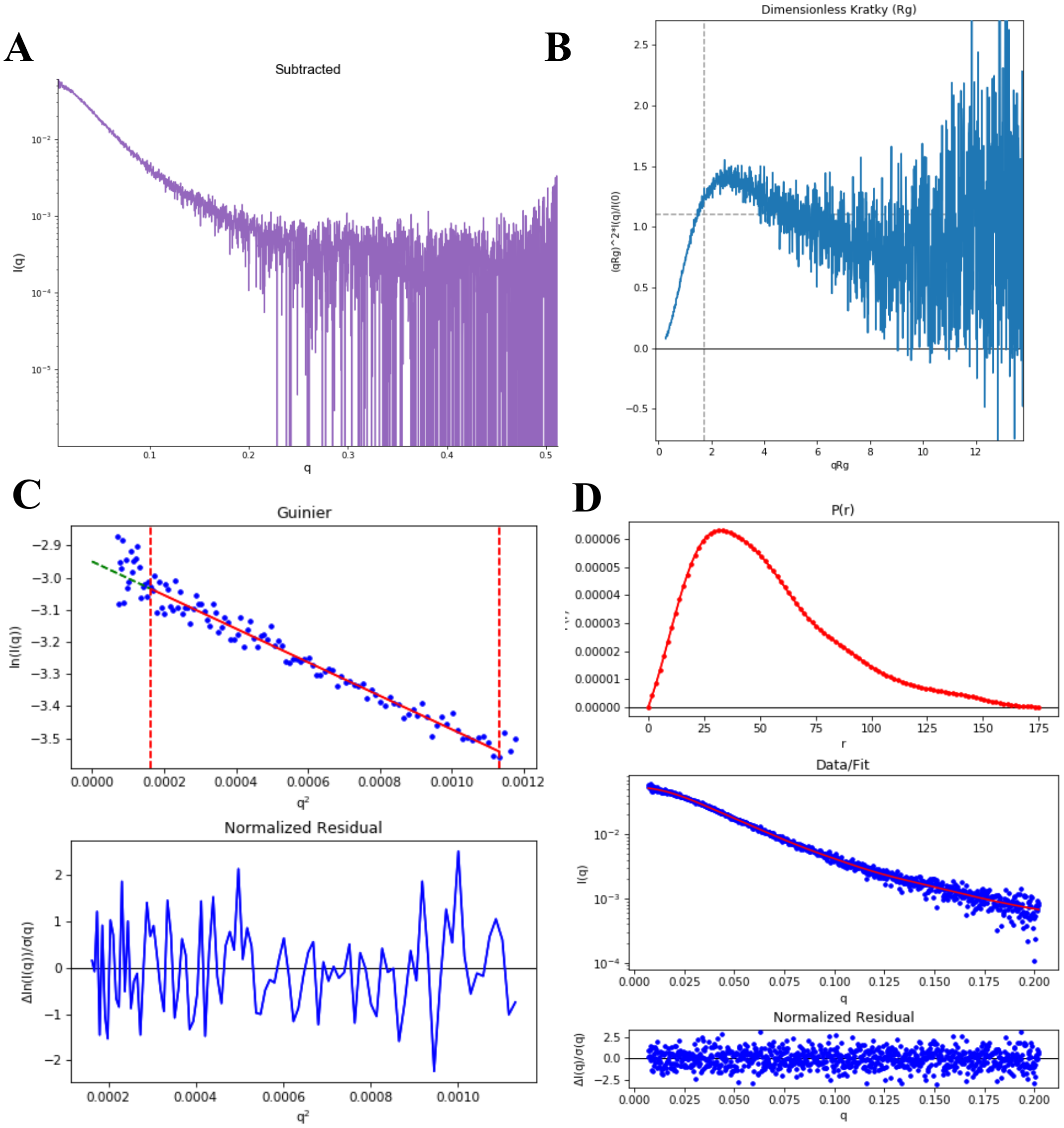


**Figure S20. SAXS plots for Nup358-KLC2/W2224A/D2225A 1.2 mg/mL.** (A) The scattering intensity profile I (q) is shown as a function of the scattering vector (q). (B) The dimensionless Kratky plot. (C) The Guinier plot (top) is shown with the normalized residual of the Guinier fit (bottom). (D) The pair distance distribution function P(r) (top), the middle panel shows the scattering intensity profile calculated from the p(r) function (red) overlaid with the scattering intensity profile of the data (blue). The bottom panel shows the normalized residual of the fit.

**Nup358-KLC2/W2224A/D2225A fusion 1 mg/mL**

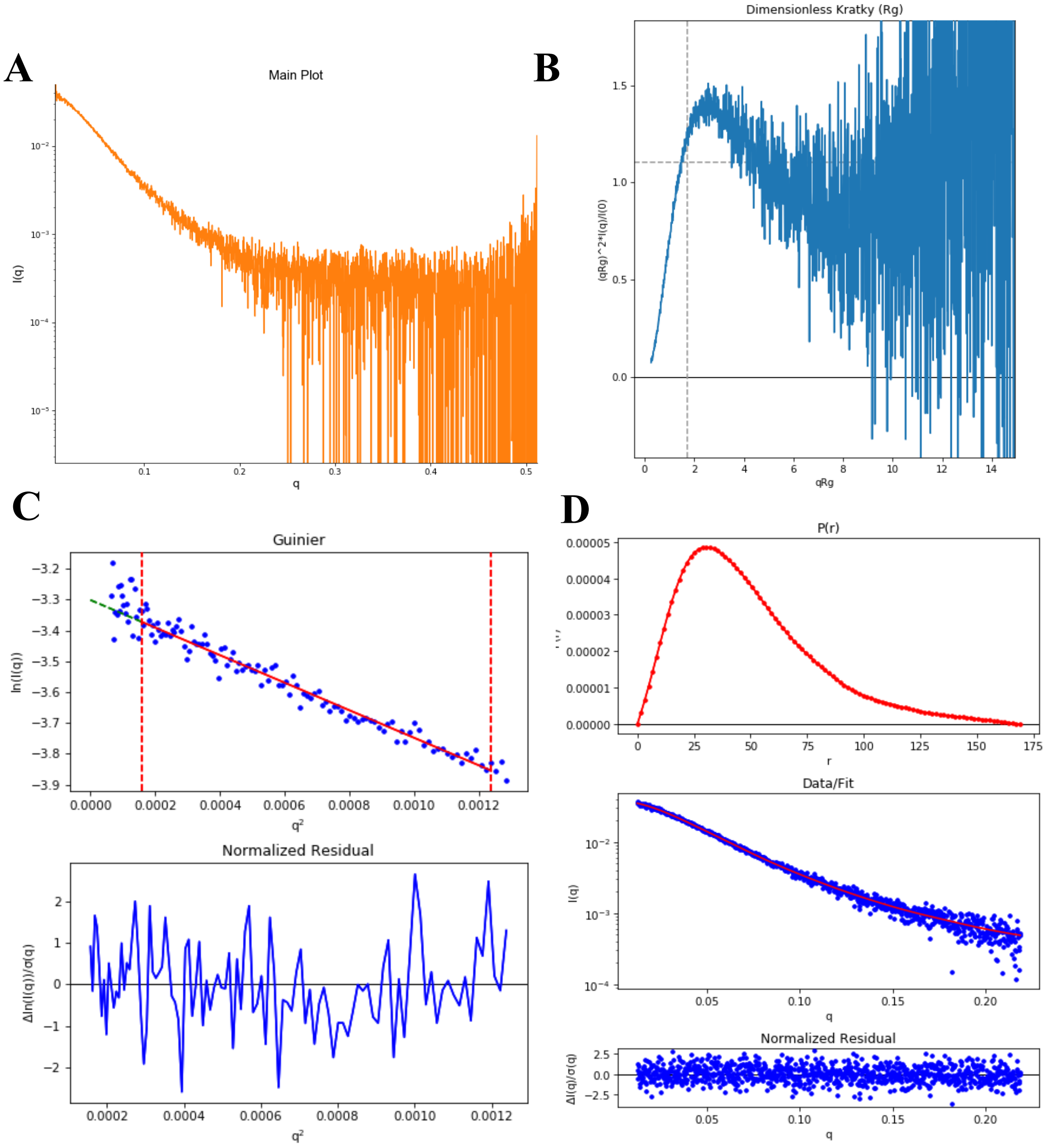


**Figure S21. SAXS plots for Nup358-KLC2/W2224A/D2225A 1 mg/mL.** (A) The scattering intensity profile I (q) is shown as a function of the scattering vector (q). (B) The dimensionless Kratky plot. (C) The Guinier plot (top) is shown with the normalized residual of the Guinier fit (bottom). (D) The pair distance distribution function P(r) (top), the middle panel shows the scattering intensity profile calculated from the p(r) function (red) overlaid with the scattering intensity profile of the data (blue). The bottom panel shows the normalized residual of the fit.

**Nup358-KLC2/W2224A/D2225A fusion 0.8 mg/mL**

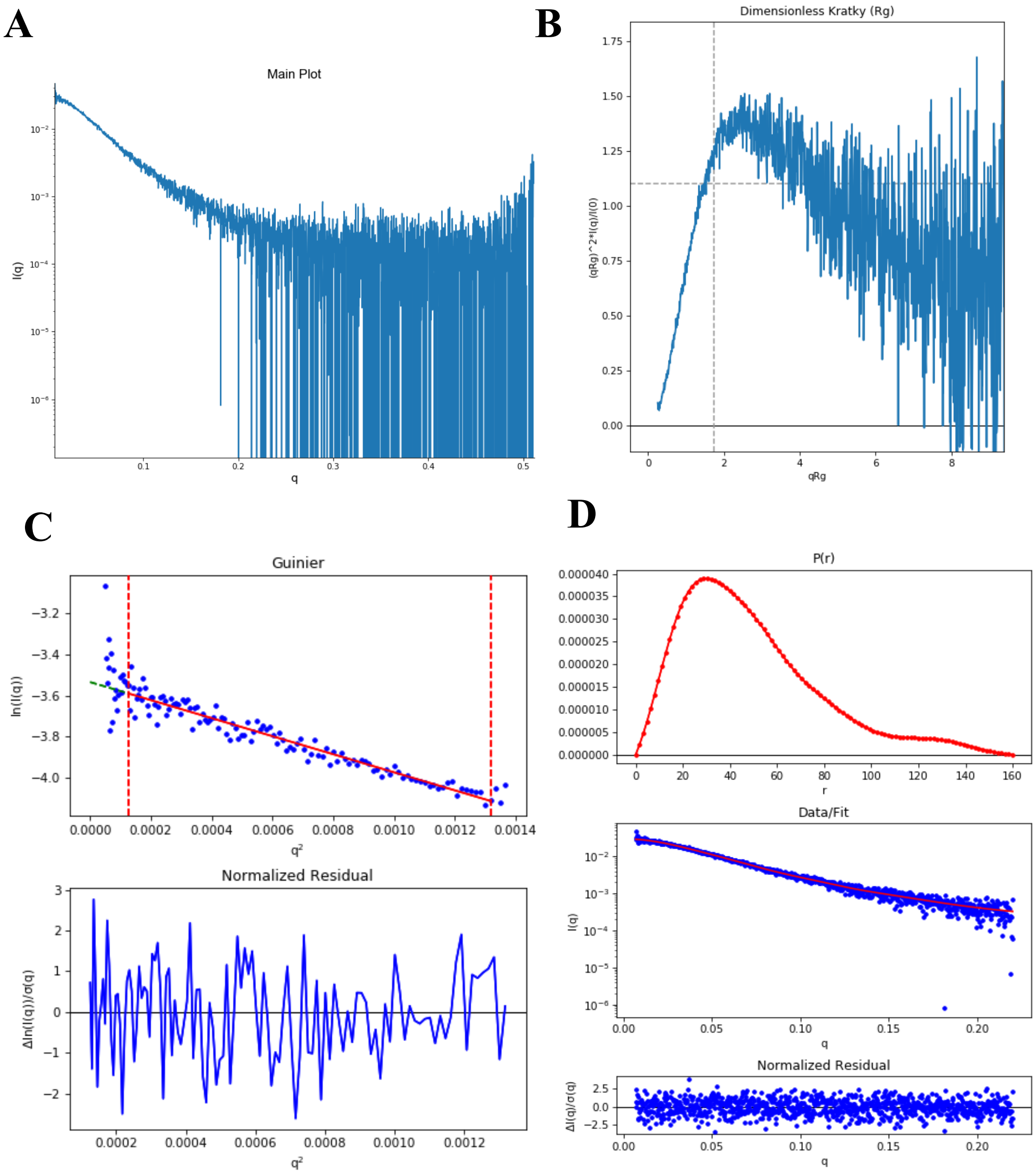


**Figure S22. SAXS plots for Nup358-KLC2/W2224A/D2225A 0.8 mg/mL.** (A) The scattering intensity profile I (q) is shown as a function of the scattering vector (q). (B) The dimensionless Kratky plot. (C) The Guinier plot (top) is shown with the normalized residual of the Guinier fit (bottom). (D) The pair distance distribution function P(r) (top), the middle panel shows the scattering intensity profile calculated from the p(r) function (red) overlaid with the scattering intensity profile of the data (blue). The bottom panel shows the normalized residual of the fit.

**Nup358-KLC2/W2224A/D2225A fusion 0.6 mg/mL**

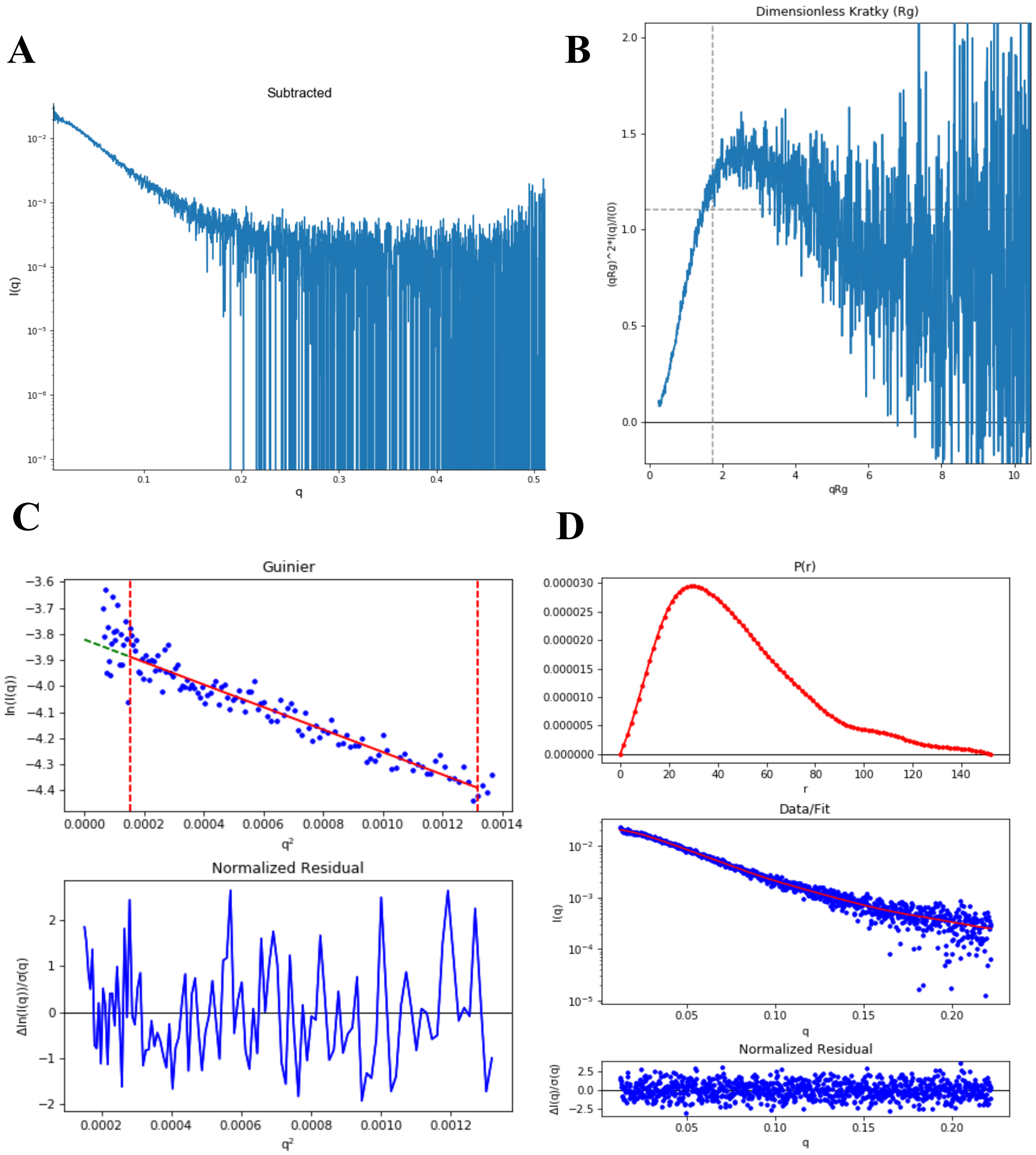


**Figure S23. SAXS plots for Nup358-KLC2/W2224A/D2225A 0.6 mg/mL.** (A) The scattering intensity profile I (q) is shown as a function of the scattering vector (q). (B) The dimensionless Kratky plot. (C) The Guinier plot (top) is shown with the normalized residual of the Guinier fit (bottom). (D) The pair distance distribution function P(r) (top), the middle panel shows the scattering intensity profile calculated from the p(r) function (red) overlaid with the scattering intensity profile of the data (blue). The bottom panel shows the normalized residual of the fit.

**Nup358-KLC2/W2224A/D2225A fusion 0.4 mg/mL**

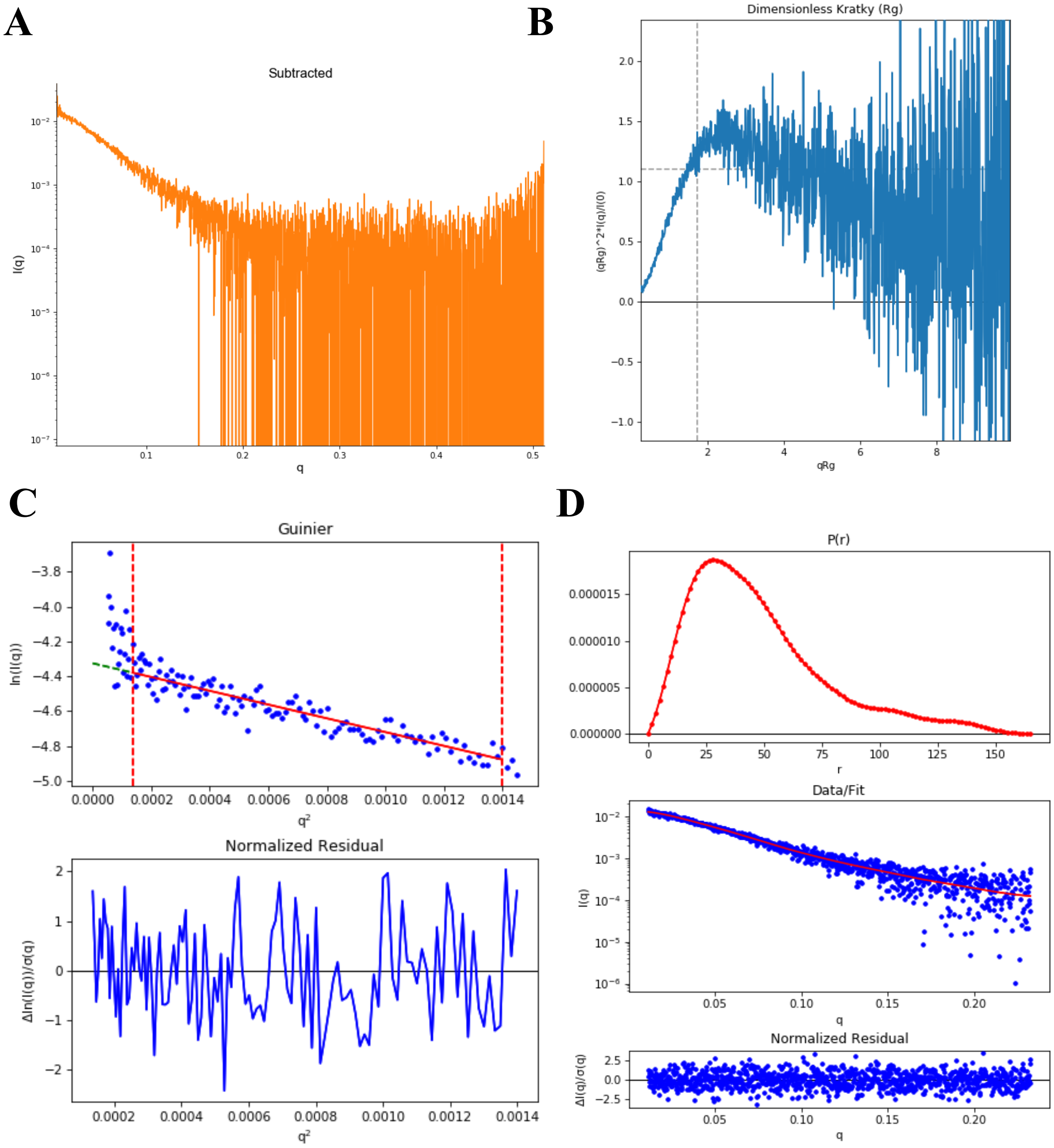


**Figure S24. SAXS plots for Nup358-KLC2/W2224A/D2225A 0.4 mg/mL.** (A) The scattering intensity profile I (q) is shown as a function of the scattering vector (q). (B) The dimensionless Kratky plot. (C) The Guinier plot (top) is shown with the normalized residual of the Guinier fit (bottom). (D) The pair distance distribution function P(r) (top), the middle panel shows the scattering intensity profile calculated from the p(r) function (red) overlaid with the scattering intensity profile of the data (blue). The bottom panel shows the normalized residual of the fit.

**Nup358-KLC2/W2224A/D2225A fusion 1.2 mg/mL + BicD2-CTD 0.3 mg/mL**

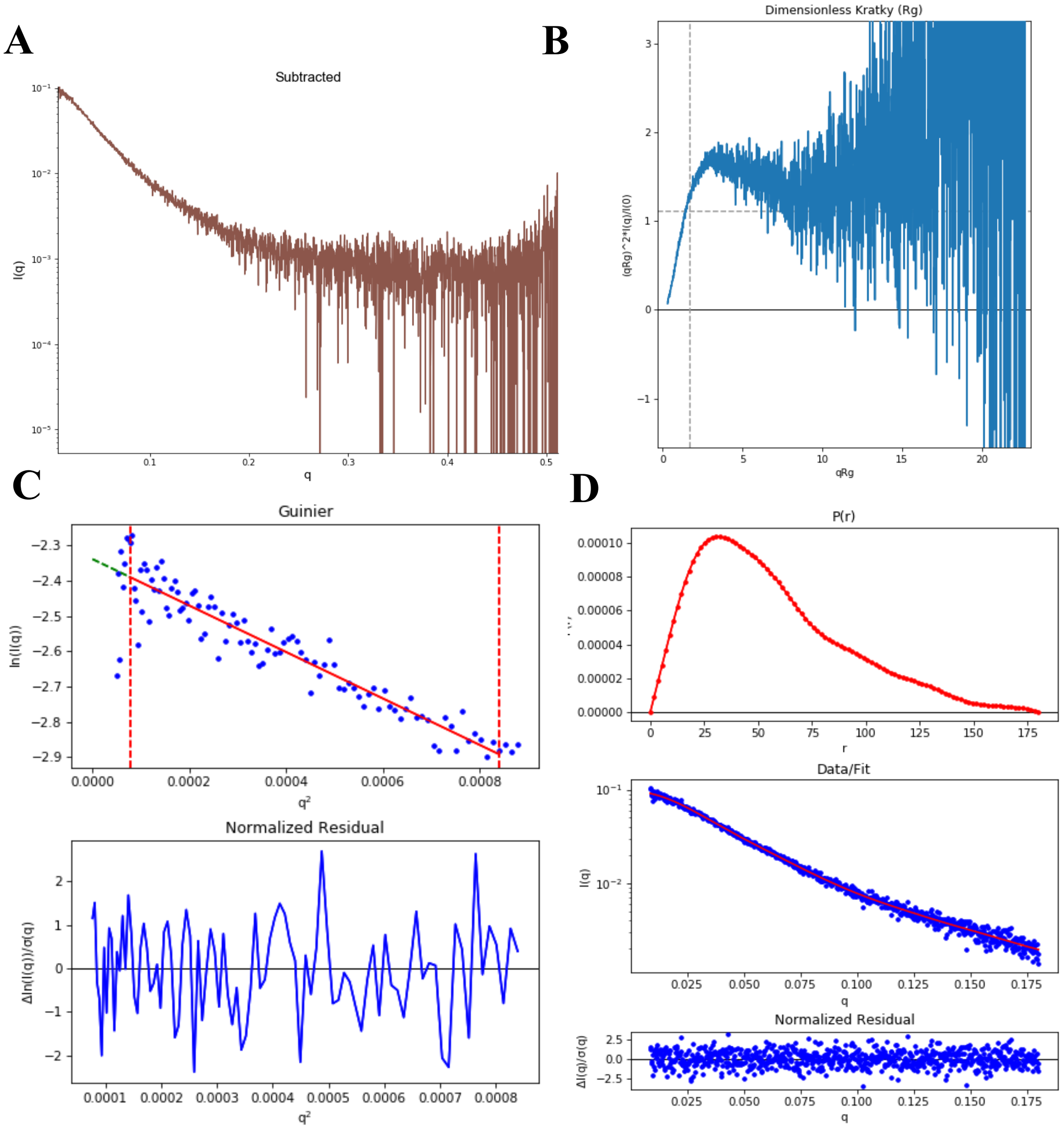


**Figure S25. SAXS plots for Nup358-KLC2/W2224A/D2225A 1.2 mg/mL+BicD2-CTD 0.3 mg/mL.** (A) The scattering intensity profile I (q) is shown as a function of the scattering vector (q). (B) The dimensionless Kratky plot. (C) The Guinier plot (top) is shown with the normalized residual of the Guinier fit (bottom). (D) The pair distance distribution function P(r) (top), the middle panel shows the scattering intensity profile calculated from the p(r) function (red) overlaid with the scattering intensity profile of the data (blue). The bottom panel shows the normalized residual of the fit.

**Nup358-KLC2/W2224A/D2225A fusion 1 mg/mL + BicD2-CTD 0.25 mg/mL**

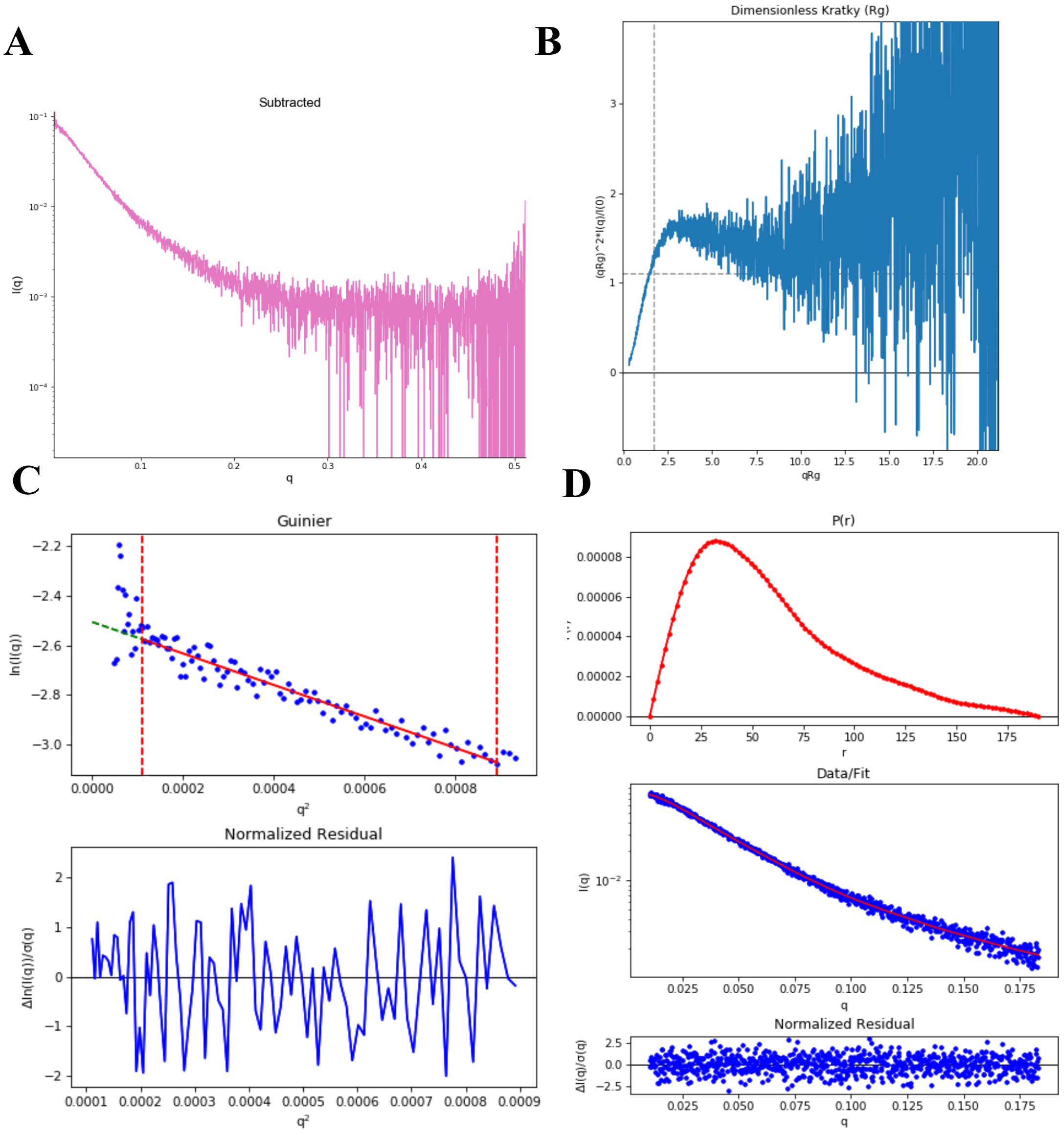


**Figure S26. SAXS plots for Nup358-KLC2/W2224A/D2225A 1 mg/mL+BicD2-CTD 0.25 mg/mL.** (A) The scattering intensity profile I (q) is shown as a function of the scattering vector (q). (B) The dimensionless Kratky plot. (C) The Guinier plot (top) is shown with the normalized residual of the Guinier fit (bottom). (D) The pair distance distribution function P(r) (top), the middle panel shows the scattering intensity profile calculated from the p(r) function (red) overlaid with the scattering intensity profile of the data (blue). The bottom panel shows the normalized residual of the fit.

**Nup358-KLC2/W2224A/D2225A fusion 0.8 mg/mL+ BicD2-CTD 0.2 mg/mL**

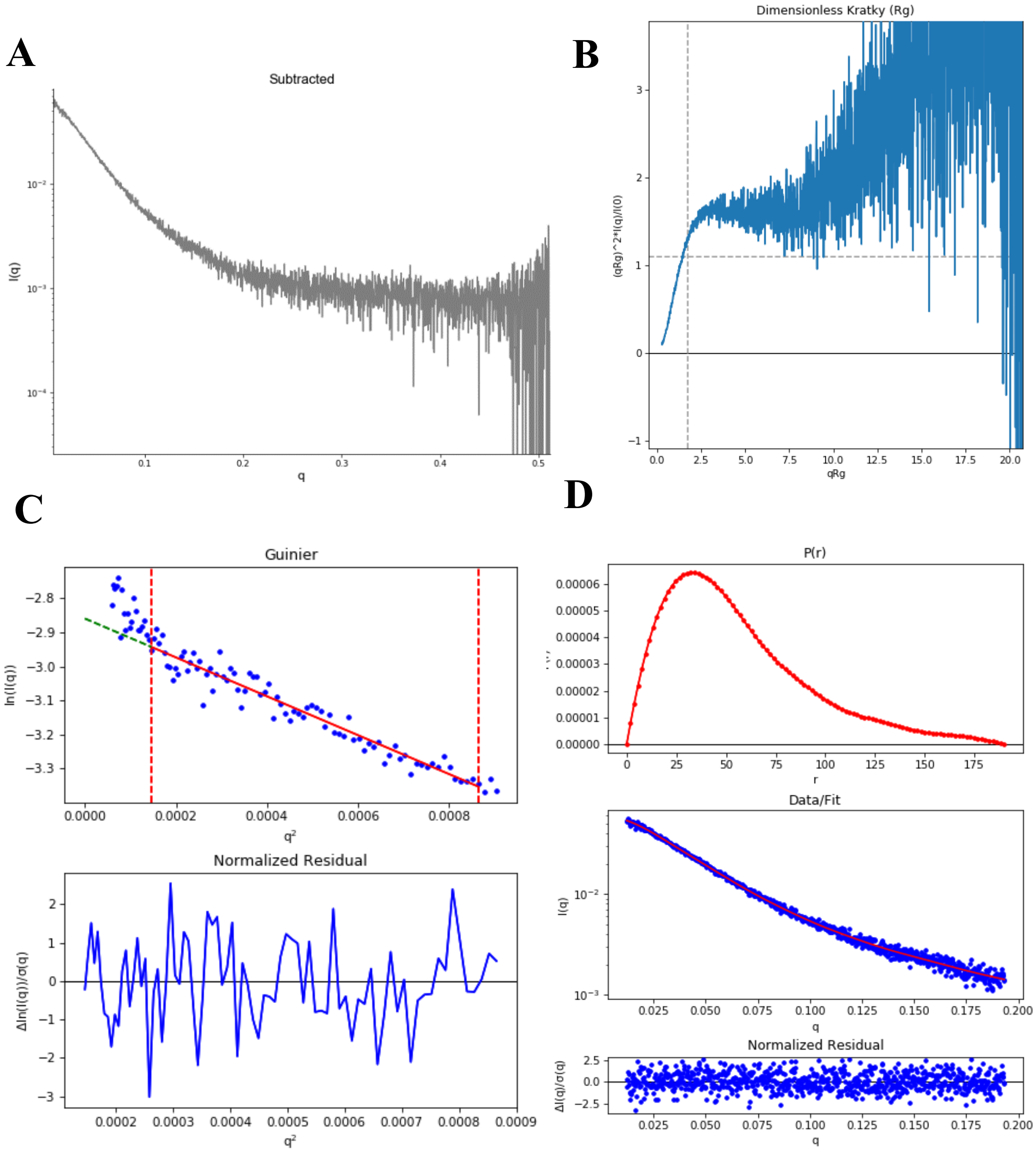


**Figure S27. SAXS plots for Nup358-KLC2/W2224A/D2225A 0.8 mg/mL+BicD2-CTD 0.2 mg/mL.** (A) The scattering intensity profile I (q) is shown as a function of the scattering vector (q). (B) The dimensionless Kratky plot. (C) The Guinier plot (top) is shown with the normalized residual of the Guinier fit (bottom). (D) The pair distance distribution function P(r) (top), the middle panel shows the scattering intensity profile calculated from the p(r) function (red) overlaid with the scattering intensity profile of the data (blue). The bottom panel shows the normalized residual of the fit.

**Nup358-KLC2/W2224A/D2225A fusion 0.6 mg/mL + BicD2-CTD 0.15 mg/mL**

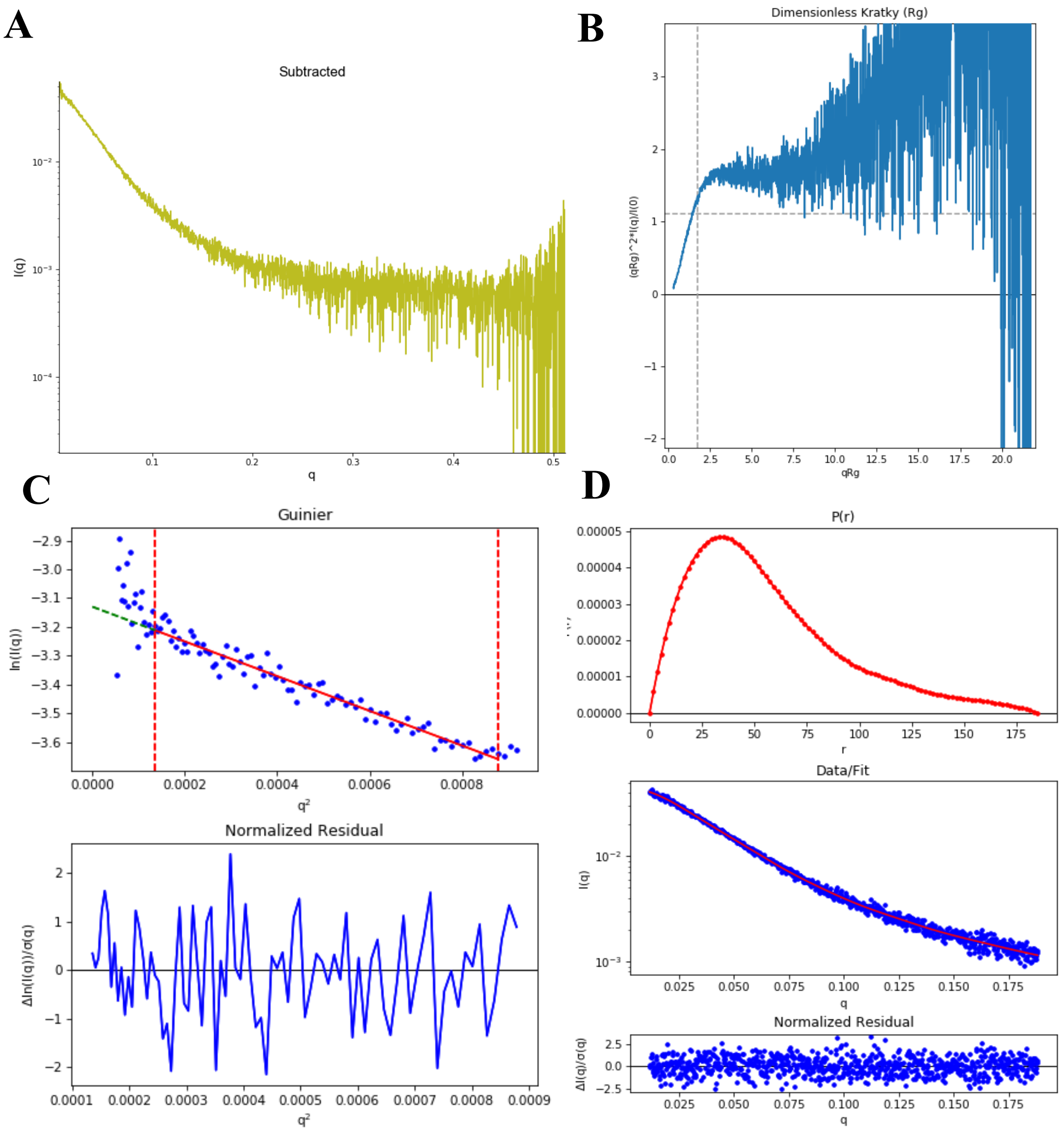


**Figure S28. SAXS plots for Nup358-KLC2/W2224A/D2225A 0.6 mg/mL+BicD2-CTD 0.15 mg/mL.** (A) The scattering intensity profile I (q) is shown as a function of the scattering vector (q). (B) The dimensionless Kratky plot. (C) The Guinier plot (top) is shown with the normalized residual of the Guinier fit (bottom). (D) The pair distance distribution function P(r) (top), the middle panel shows the scattering intensity profile calculated from the p(r) function (red) overlaid with the scattering intensity profile of the data (blue). The bottom panel shows the normalized residual of the fit.

**Nup358-KLC2/W2224A/D2225A fusion 0.4 mg/mL + BicD2-CTD 0.1 mg/mL**

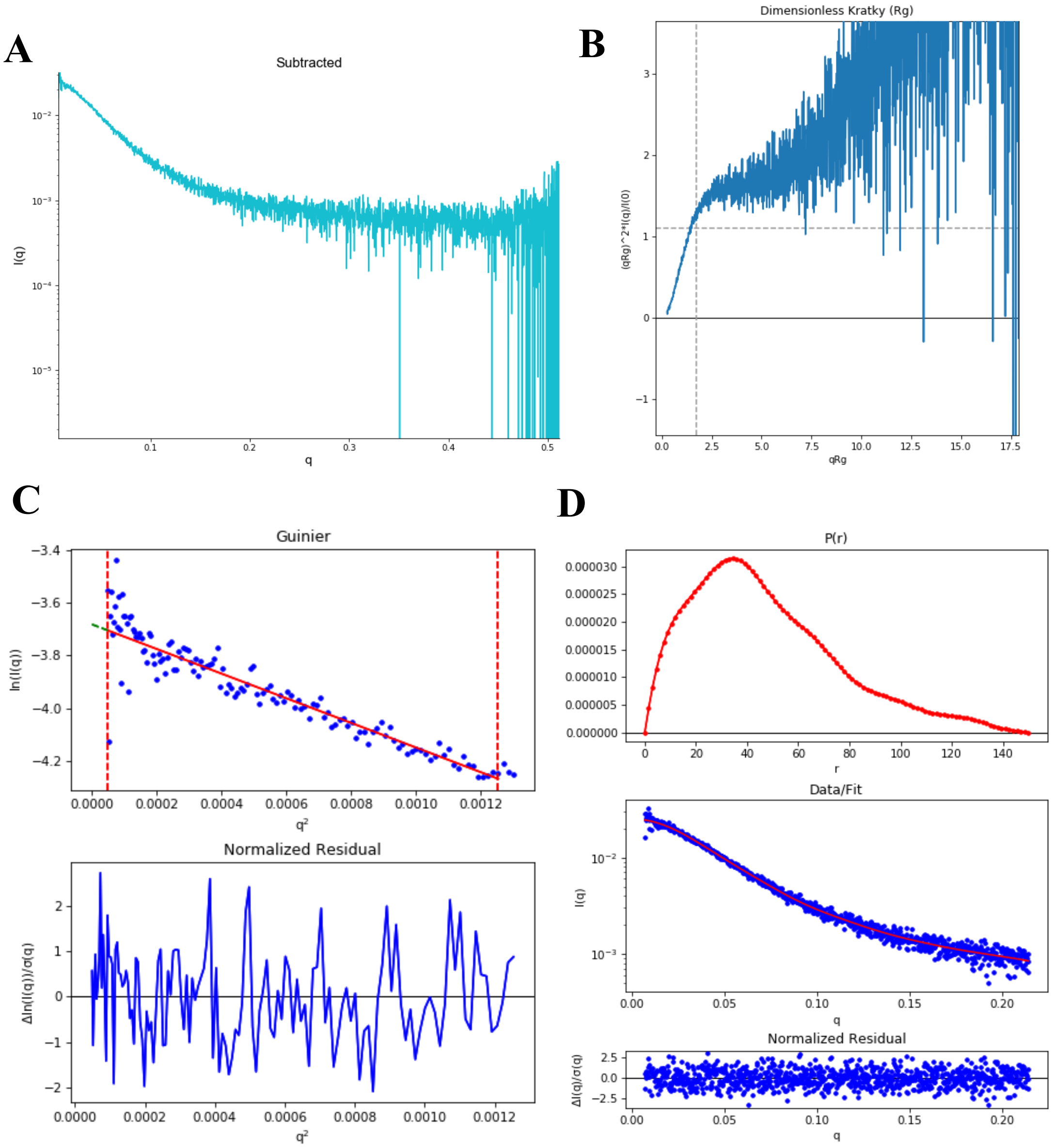


**Figure S29. SAXS plots for Nup358-KLC2/W2224A/D2225A 0.4 mg/mL+BicD2-CTD 0.1 mg/mL.** (A) The scattering intensity profile I (q) is shown as a function of the scattering vector (q). (B) The dimensionless Kratky plot. (C) The Guinier plot (top) is shown with the normalized residual of the Guinier fit (bottom). (D) The pair distance distribution function P(r) (top), the middle panel shows the scattering intensity profile calculated from the p(r) function (red) overlaid with the scattering intensity profile of the data (blue). The bottom panel shows the normalized residual of the fit.

**Nup358-KLC2/W2224A/D2225A fusion 0.2 mg/mL + BicD2-CTD 0.05 mg/mL**

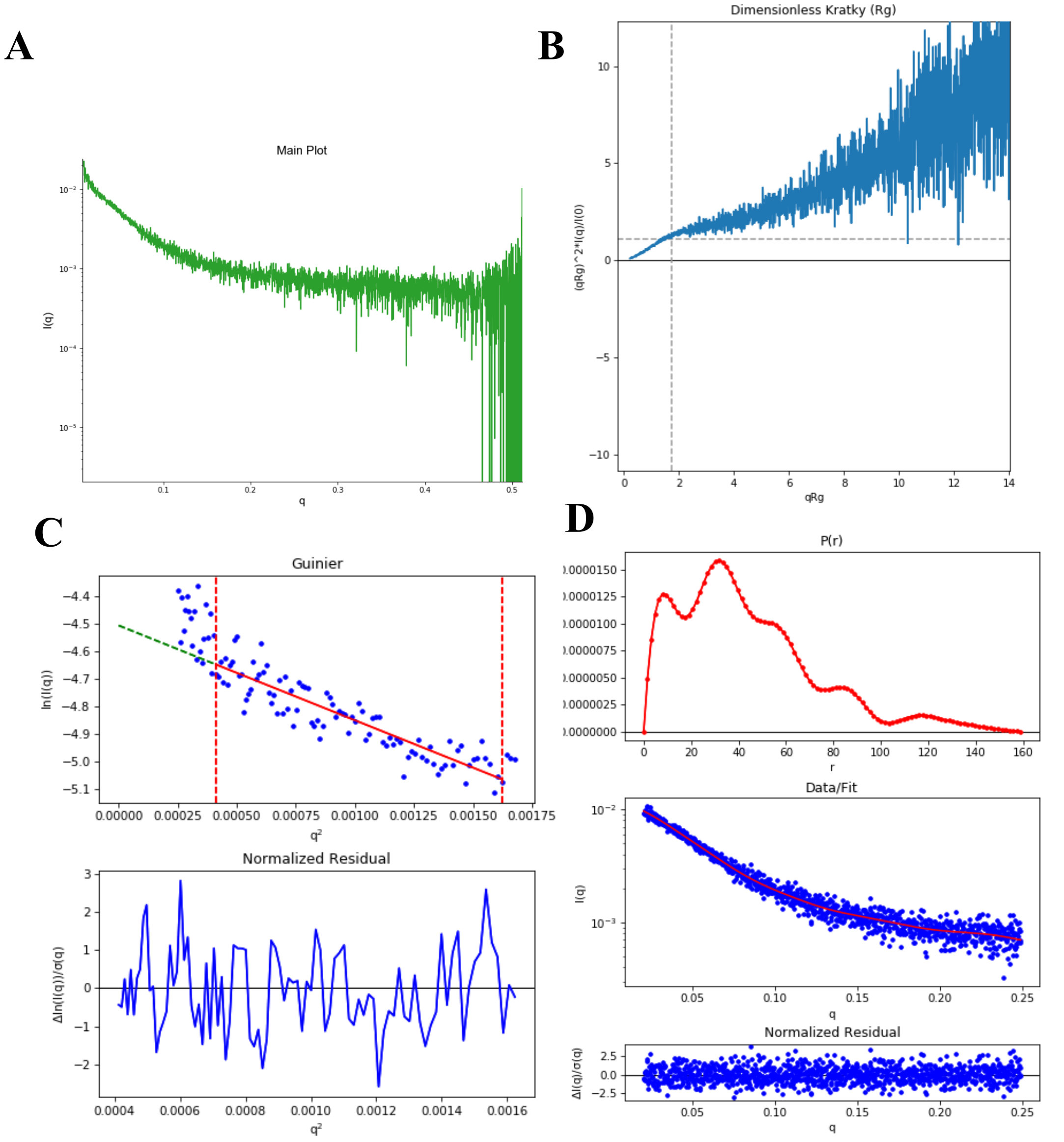


**Figure S30. SAXS plots for Nup358-KLC2/W2224A/D2225A 0.2 mg/mL+BicD2-CTD 0.05 mg/mL.** (A) The scattering intensity profile I (q) is shown as a function of the scattering vector (q). (B) The dimensionless Kratky plot. (C) The Guinier plot (top) is shown with the normalized residual of the Guinier fit (bottom). (D) The pair distance distribution function P(r) (top), the middle panel shows the scattering intensity profile calculated from the p(r) function (red) overlaid with the scattering intensity profile of the data (blue). The bottom panel shows the normalized residual of the fit. Note that this analysis is poor due to low signal and high noise due to the sample having very low concentration and is only included for comparison, data was not used for further analysis.
